# Protein-mediated genome folding allosterically enhances site-specific integration of foreign DNA into CRISPRs

**DOI:** 10.1101/2023.05.26.542337

**Authors:** Andrew Santiago-Frangos, William S. Henriques, Tanner Wiegand, Colin C. Gauvin, Murat Buyukyoruk, Ava B. Graham, Royce A. Wilkinson, Lenny Triem, Kasahun Neselu, Edward T. Eng, Gabriel C. Lander, Blake Wiedenheft

**Affiliations:** Department of Microbiology and Cell Biology, Montana State University, Bozeman, MT, USA; Department of Chemistry and Biochemistry, Montana State University, Bozeman, MT, USA; Thermal Biology Institute, Montana State University, Bozeman, MT, USA; Simons Electron Microscopy Center, National Resource for Automated Molecular Microscopy, New York Structural Biology Center, New York, NY, USA; Department of Integrative Structural and Computational Biology, The Scripps Research Institute, La Jolla, CA, USA

**Keywords:** CRISPR-Cas, integration, CRISPR adaptation, Cas1-2/3, IHF, DNA recording devices, transposition, transposase, DNA bending

## Abstract

Bacteria and archaea acquire resistance to viruses and plasmids by integrating fragments of foreign DNA into the first repeat of a CRISPR array. However, the mechanism of site-specific integration remains poorly understood. Here, we determine a 560 kDa integration complex structure that explains how Cas (Cas1-2/3) and non-Cas proteins (IHF) fold 150 base-pairs of host DNA into a U-shaped bend and a loop that protrude from Cas1-2/3 at right angles. The U-shaped bend traps foreign DNA on one face of the Cas1-2/3 integrase, while the loop places the first CRISPR repeat in the Cas1 active site. Both Cas3s rotate 100-degrees to expose DNA binding sites on either side of the Cas2 homodimer, that each bind an inverted repeat motif in the leader. Leader sequence motifs direct Cas1-2/3-mediated integration to diverse repeat sequences that have a 5’-GT.

---

Vertebrates, bacteria, and archaea have domesticated transposases (e.g., RAG1 and Cas1) for adaptive immunity ^1,2^. Integrases, transposases and recombinases often co-opt additional DNA-bending proteins (e.g., IHF, HU, H-NS, or HMGB1) that facilitate DNA integration and excision ^3–7^. However, the structural role of DNA folding during this mobilization of DNA remains largely enigmatic.

<u>C</u>lustered <u>R</u>egularly Interspaced <u>S</u>hort <u>P</u>alindromic <u>R</u>epeats (CRISPRs) are essential components of an adaptive immune system that stores DNA-based molecular memories of past infections ^8^. <u>C</u>RISPR-<u>as</u>sociated proteins, Cas1 and Cas2, integrate fragments of foreign DNA ("spacers") into CRISPRs. Integration duplicates a repeat sequence, which thereby maintains the characteristic repeat-spacer-repeat architecture (**Fig. 1a**). Cas1 and Cas2 form a heterohexameric complex that consists of two Cas1 homodimers (Cas1a-a* and Cas1b-b*) flanking a Cas2 homodimer (**Fig. 1a-b**) ^9–11^. Foreign DNA fragments bind across one face of the Cas2 homodimer, which positions the 3’-ends into Cas1 active sites on either end of the complex (i.e., Cas1a* and Cas1b*) ^8–10,12^. The CRISPR repeat sequence wraps around the opposing face of Cas2, sandwiching the Cas2 homodimer between the foreign and repeat DNA duplexes. Opposing Cas1 subunits (Cas1a* and Cas1b*) catalyze two successive strand-transfer reactions, linking the 3’-ends of the foreign DNA to opposite ends of the repeat ^5,11^. CRISPR integration complexes sense a 2-5 bp 3’ overhang called a protospacer-adjacent motif (PAM) in the foreign DNA to determine the integration orientation ^9^. Correct spacer orientation is necessary to produce a functional CRISPR RNA that guides the CRISPR interference machinery (i.e., Cascade) to complementary targets ^8,13^. Integration occurs in a stepwise manner. First, the non-PAM end of the foreign DNA is integrated at the leader-side of the repeat ^14^. Second, the PAM is cleaved by Cas or non-Cas nucleases and the trimmed 3’ end is integrated at the spacer-side of the repeat ^14–16^. These integration events tie a non-covalent knot around the Cas2 homodimer (foreign DNA on one side and repeat DNA on the other) that is held together by complementary base-pairing in the foreign DNA.

**Figure 1.**
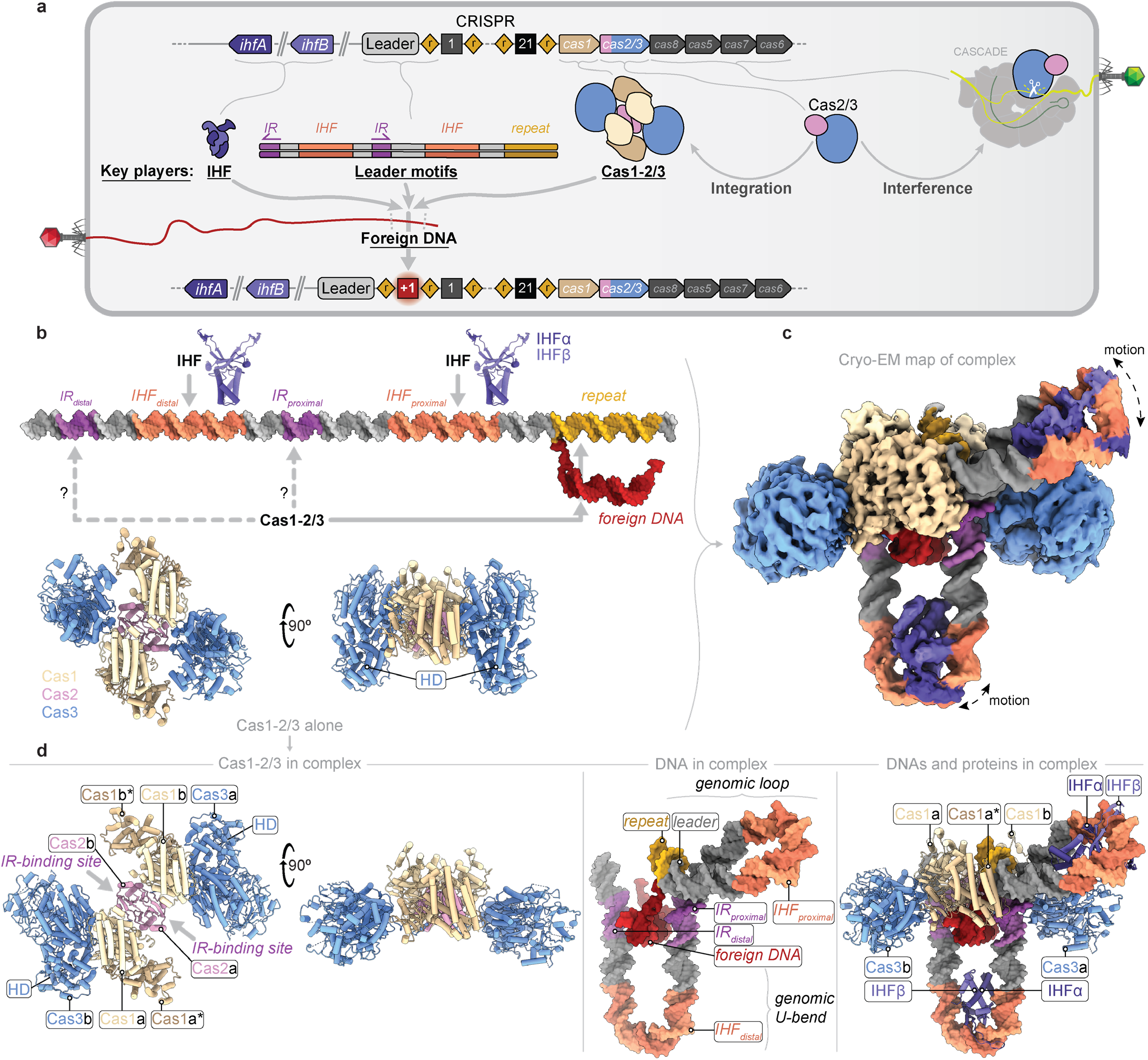
Cryo-EM structure of the type I-F CRISPR integration complex. **a**, Scheme of the I-F CRISPR system of *Pseudomonas aeruginosa* PA14. A CRISPR is composed of repeated DNA sequences (diamonds) interspersed with unique spacer sequences (black squares). The CRISPR is adjacent to six *cas* genes (arrows). Four Cas1 and two Cas2/3 proteins assemble into a heterohexamer in which the Cas3 and Cas1 subunits surround the central Cas2 homodimer like petals of a closed flower (Cas14-Cas2/32). Cas1-2/3 and integration host factor (IHF) proteins cooperate with DNA upstream of the CRISPR (leader) to integrate foreign DNA at the first repeat. The leader sequence contains two IHF binding sites and two Inverted Repeats (IRs) that are necessary for integration of foreign DNA at the leader-repeat junction. In addition to playing a central role in integration, the Cas2/3 fusion is recruited to DNA-bound Cascade CRISPR surveillance complex to degrade foreign genetic parasites. **b**, The Cas1-2/3 heterohexamer and IHF proteins were mixed with a half-site DNA integration intermediate consisting of a foreign DNA linked to one strand of the CRISPR DNA at the leader-repeat junction. **c**, Cryo-EM density map of the type I-F CRISPR integration complex at ∼3.5 Å resolution (**Extended Data Fig. 1g-k, Table 1**). **d**, Atomic model of the type I-F CRISPR integration complex. Cas1-2/3 proteins alone (left) are shown in cartoon representation. Cas3 domains rotate by 100° simulating the motion of a bloomed flower and exposing DNA binding sites on Cas2 that interact with each of the IRs (**Extended Data Fig. 2a and Supplementary Video 1**). DNA alone is shown in the middle (surface representation). Proteins (cartoon representations) and DNAs of the integration complex are shown on the right.

**Table 1.**
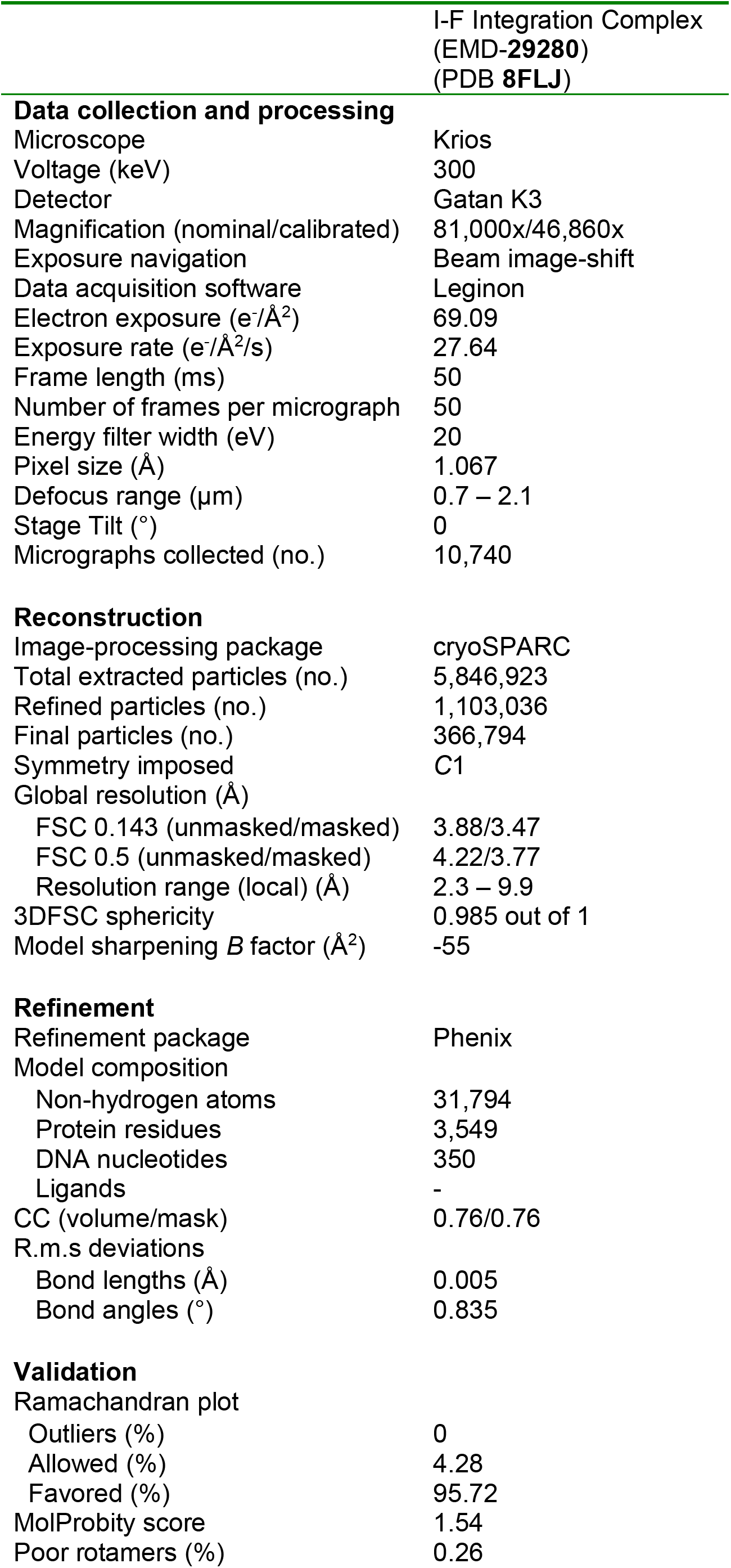
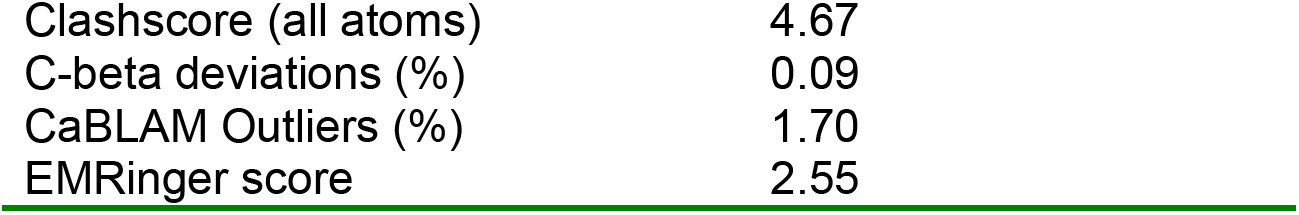
Cryo-EM data collection, refinement, and validation statistics.

|  | I-F Integration Complex<br>(EMD-29280)<br>(PDB 8FLJ) |
| --- | --- |
| **Data collection and processing** |  |
| Microscope | Krios |
| Voltage (keV) | 300 |
| Detector | Gatan K3 |
| Magnification (nominal/calibrated) | 81,000x/46,860x |
| Exposure navigation | Beam image-shift |
| Data acquisition software | Leginon |
| Electron exposure (e<sup>-</sup>/Å<sup>2</sup>) | 69.09 |
| Exposure rate (e<sup>-</sup>/Å<sup>2</sup>/s) | 27.64 |
| Frame length (ms) | 50 |
| Number of frames per micrograph | 50 |
| Energy filter width (eV) | 20 |
| Pixel size (Å) | 1.067 |
| Defocus range (μm) | 0.7 – 2.1 |
| Stage Tilt (°) | 0 |
| Micrographs collected (no.) | 10,740 |
| **Reconstruction** |  |
| Image-processing package | cryoSPARC |
| Total extracted particles (no.) | 5,846,923 |
| Refined particles (no.) | 1,103,036 |
| Final particles (no.) | 366,794 |
| Symmetry imposed | C1 |
| Global resolution (Å) |  |
| FSC 0.143 (unmasked/masked) | 3.88/3.47 |
| FSC 0.5 (unmasked/masked) | 4.22/3.77 |
| Resolution range (local) (Å) | 2.3 – 9.9 |
| 3DFSC sphericity | 0.985 out of 1 |
| Model sharpening *B* factor (Å<sup>2</sup>) | -55 |
| **Refinement** |  |
| Refinement package | Phenix |
| Model composition |  |
| Non-hydrogen atoms | 31,794 |
| Protein residues | 3,549 |
| DNA nucleotides | 350 |
| Ligands | - |
| CC (volume/mask) | 0.76/0.76 |
| R.m.s deviations |  |
| Bond lengths (Å) | 0.005 |
| Bond angles (°) | 0.835 |
| **Validation** |  |
| Ramachandran plot |  |
| Outliers (%) | 0 |
| Allowed (%) | 4.28 |
| Favored (%) | 95.72 |
| MolProbity score | 1.54 |
| Poor rotamers (%) | 0.26 |
| Clashscore (all atoms) | 4.67 |
| C-beta deviations (%) | 0.09 |
| CaBLAM Outliers (%) | 1.70 |
| EMRinger score | 2.55 |

New foreign DNA is preferentially integrated at the first repeat in a CRISPR locus, ensuring efficient transcription and processing of CRISPR RNAs that target the most recently encountered genetic parasites (**Fig. 1a**) ^17,18^. Cas1-2 is thought to recognize a palindromic sequence within the CRISPR repeat, similar to target site recognition by many DNA transposases ^5,11,19–21^. However, Cas1-2 recognition of the palidromic repeat does not explain how the first repeat is differentiated from downstream repeat sequences in a CRISPR. Thus, polarized integration often relies on additional proteins and DNA sequence motifs upstream of the CRISPR (i.e., leader) ^3–5,11,18,22–27^. Integration <u>H</u>ost <u>F</u>actor (IHF) facilitates polarized integration in the type I-E CRISPR system from *Escherichia coli* ^3^. A structure of the I-E integration complex revealed that IHF bends the leader DNA to bring an upstream sequence motif into contact with Cas1, and IHF further stabilizes the Cas1-2 integrase at the first repeat through direct Cas1-IHF interactions ^3,5^.

Cas1 and Cas2 are conserved components of CRISPR-mediated immune systems. However, the type I-F CRISPR system has a unique fusion of the Cas2 subunit to the Cas3-nuclease/helicase found in many type I systems (**Fig. 1a-b**) ^28,29^. Cas3 degrades Cascade-bound DNA into fragments with PAM-containing termini, that are captured by Cas1-2 and integrated into the CRISPR locus, in a process called “primed acquisition” ^4,30–37^. However, the structural mechanism for primed acquisition is unclear. Additionally, in contrast to the *E. coli* I-E system, which relies on one IHF to bend DNA and recruit a DNA motif found ∼50 bp upstream of the CRISPR repeat, most I-F, I-C, and some I-E CRISPR leaders, contain multiple IHF binding sites and multiple subtype-specific DNA motifs found up to 100-200 bp upstream of the CRISPR repeat (**Fig. 1a and Extended Data Fig. 1a**) ^22^.

To determine how DNA sequence motifs in the CRISPR leader regulate the Cas1-2/3 integrase, we determined the structure of a ∼560 kDa CRISPR integration complex from *Pseudomonas aeruginosa*. The structure reveals that Cas1-2/3 cannot interact with the leader without first undergoing a large conformational change that may be induced by foreign DNA binding. Further, the structure explains how the I-F leader and IHF proteins guide Cas1-2/3 to deliver and integrate foreign DNA at the first repeat in the CRISPR array (**Fig. 1**). Cas1-2/3 and IHF interact with all five DNA sequence motifs (i.e., two Inverted Repeats, two IHF binding sites, and the CRISPR repeat) primarily through a shape-based readout ^38,39^. The shape of the folded I-F CRISPR leader is similar to that of the λ-phage excision complex, suggesting that DNA is often used as a flexible scaffold to regulate DNA mobilization ^3–7,40–53^. The structure suggests that site-specific integration relies on protein-induced folding of the upstream DNA rather than sequence-specific recognition of the repeat. To test this idea, we perform a series of integration reactions demonstrating that efficient integration relies on conserved sequences in the leader and a 5’ GT dinucleotide in the repeat. We show that 5’ GT dinucleotides are broadly conserved in repeats derived from different CRISPR types, suggesting that they play a conserved role in integration across diverse CRISPR systems. In addition, the I-F CRISPR integration complex suggests a structural mechanism for interactions of the Cas1-2/3 integrase with the Cascade surveillance complex, that may be neccesary for rapid adaptation to phage escape mutants^4,30–37^.

## Cryo-EM structure of type I-F CRISPR integration complex

To understand how the Cas1-2/3 integrase cooperates with IHF and CRISPR leader motifs to integrate foreign DNA at the first CRISPR repeat, we purified the heterohexameric Cas1-2/3 integrase and the IHFα-β heterodimer, incubated these proteins with a DNA substrate representing a half-site integration intermediate, and isolated the assembled complex using <u>S</u>ize <u>E</u>xclusion <u>C</u>hromatography (SEC) (**Extended Data Fig. 1a-f**). The purified I-F integration complex was applied to cryo-EM grids and vitrified. We recorded 10,740 movies and picked 366,794 particles to determine a ∼3.48 Å-resolution structure of the integration complex. The reconstructed density was of sufficient to model 90.7% of the 10 polypeptides and 88.4% of the 396 nucleotides of DNA (**Fig. 1b-d, Extended Data Fig. 1g-k, Table 1**). The model explains how the Cas1-2/3 subunits cooperate with two IHF heterodimers to kink and twist ∼150 base-pairs of host DNA into a structure that precisely positions foreign DNA for integration at the first repeat of the CRISPR (**Fig. 1b-d**).

The Cas1 and Cas2 subunits adopt a familiar quaternary arrangement that binds a foreign DNA on one face of the Cas2 homodimer and CRISPR repeat DNA on the other face (**Fig. 1d, 2a and 3a**) ^8^. A previously determined structure of Cas1-2/3 alone revealed that the Cas3 and Cas1 domains surround the central Cas2 homodimer like petals of a closed flower (Cas1_4_:Cas2/3_2_) (**Fig. 1b**) ^54^. While this structure explained how Cas1 regulates the Cas3 nuclease, the role of Cas3 during integration remained unclear ^54^. Here, we show that the addition of DNA drives a series of conformational changes in both the DNA and proteins. The Cas3 domains rotate ∼100° to align in a planar configuration with Cas2, simulating the motion of a bloomed flower, and exposing equivalent surfaces on opposite sides of the Cas2 homodimer that recognize an inverted repeat (IR) that is conserved in I-F leaders (**Fig. 1d, Extended Data Fig. 2a and Supplementary Video 1**) ^54^. Thus, the new planar conformation of Cas1-2/3 enables the simultaneous coordination of four DNA helices (IR_distal_, IR_proximal_, foreign DNA, and CRISPR repeat) around the central Cas2 homodimer (**Fig. 1d and 3**). Further, this Cas3 rotation flips the nuclease domain from an interaction with Cas1 that suppresses the Cas3 nuclease activity, to the opposite side of the complex, where the back of the Cas3 nuclease domain docks onto a groove created at the Cas1-Cas1 interaface (**Fig. 1b,d and Extended Data Fig. 2a,b**).

**Figure 2.**
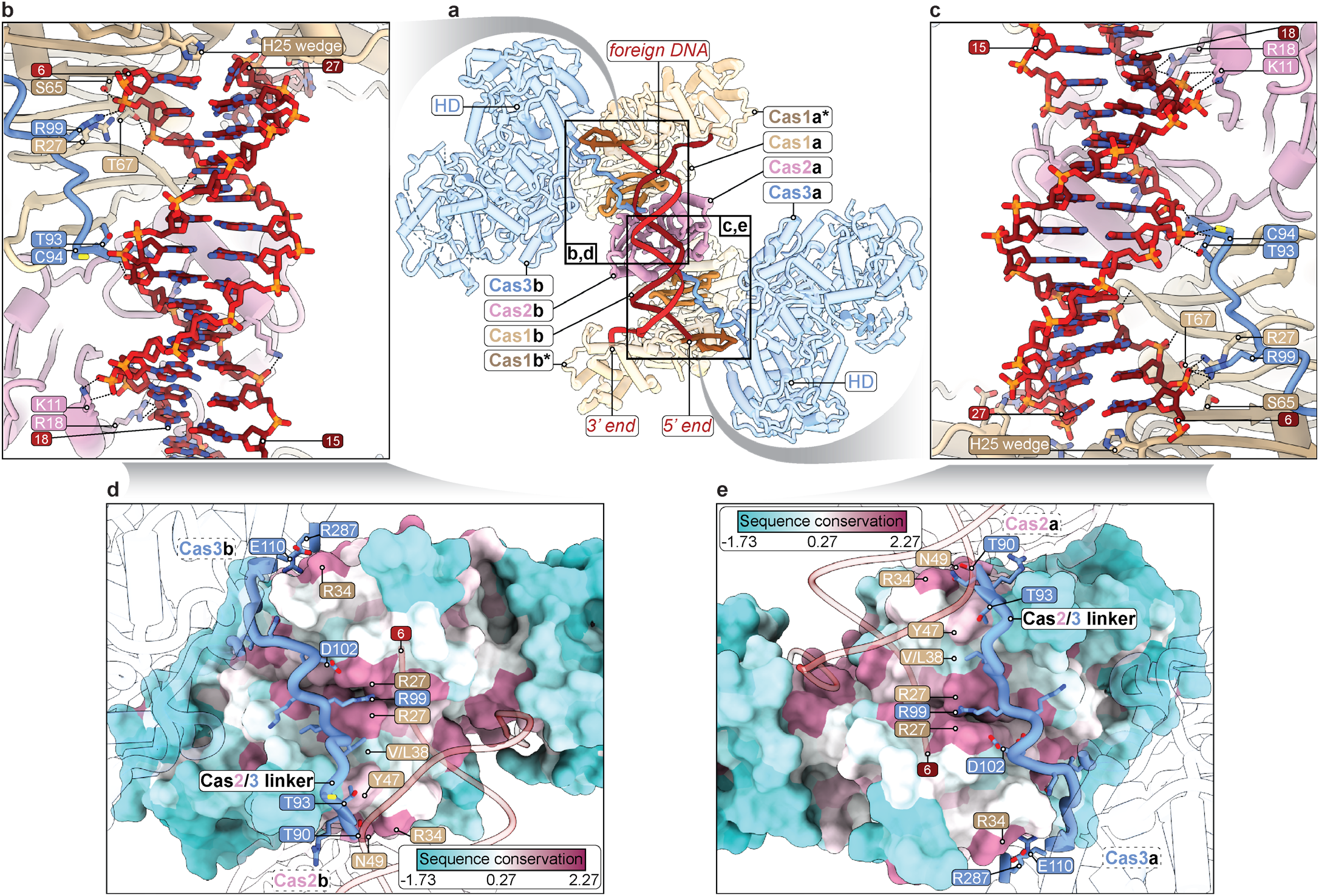
Foreign DNA constrains the Cas2/3 linker against conserved Cas1 residues. **a**, View of the foreign DNA-bound face of Cas1-2/3. The foreign DNA, Cas2 subunits, Cas2/3 linker and Cas1 beta hairpins that contact the start and end of the Cas2/3 linker are shown in solid while other parts of the complex are shown at 40% transparency for clarity. Insets outline locations of close-up views shown in panels b-e. **b**,**c**, The foreign DNA constrains the Cas2/3 linker against each Cas1 subunit (**Extended Data Fig. 2b**). Cas2, the Cas2/3 linker and, Cas1 cooperate to bind the foreign DNA body and to splay the ends of the foreign DNA. Histidine wedges in Cas1 measure out a central foreign DNA duplex of 22 base-pairs. Most DNA-binding residues are conserved or undergo conservative mutations (**Extended Data Fig. 3**). **d**,**e**, Conserved Cas2/3 linker residues (blue, sticks) contact residues conserved in Cas1 proteins from type I-F CRISPR systems (mauve, surface) (**Extended Data Fig. 2b,c**). Cas2 and Cas3 domains are shown at 90% transparency for clarity. Inset shows the Cas1 sequence conservation color key.

**Figure 3.**
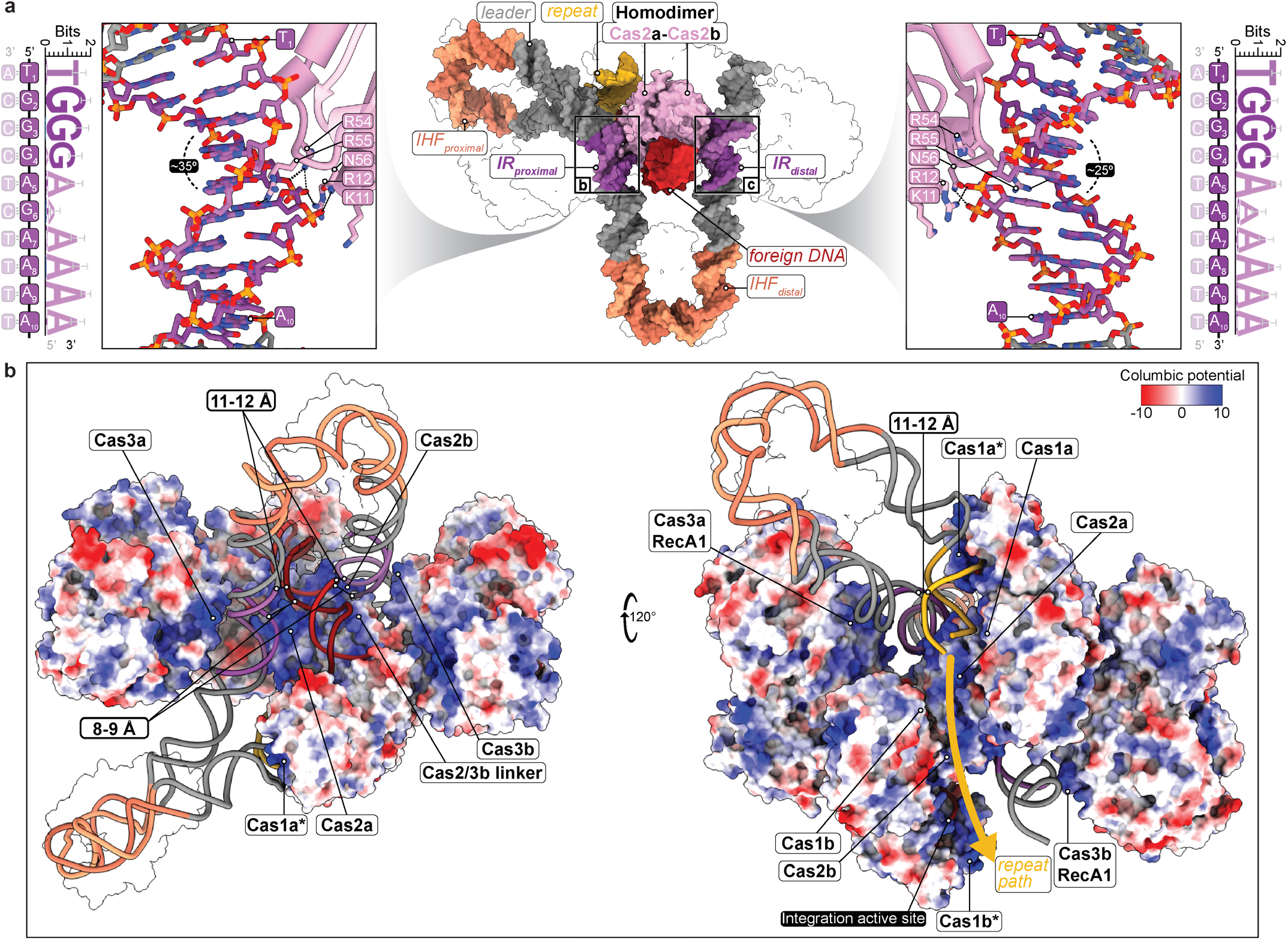
The Cas2 homodimer simultaneously coordinates four dsDNA helices critical to CRISPR integration. **a**, The Cas2 homodimer (pink surface) is flanked by DNA on four sides. Prior structures have shown that the CRISPR repeat (yellow) and foreign DNA (red) are bound to opposite faces of Cas2. Here, we show that symmetrical surfaces on Cas2 also bind inverted repeat (IR, left and right) motifs in the leader. Surface representations of the IHF heterodimers, Cas1 homodimers, Cas2/3 linkers, Cas3 domains, and the 3’ overhang of the foreign DNA are shown in 100% transparency for clarity. Each Cas2 inserts an arginine (R55) into the center of the IRs, which stack between deoxyribose sugars, and additional polar residues (R54, N56, R12, and K11) contact the DNA backbone. Cas2 induces 25-35° bends in the DNA (**Extended Data Fig. 4c,e**). The sequence logos of the type I-F IRproximal (left) and IRdistal motifs (right), and the IR sequences present in the *P. aeruginosa* PA14 CRISPR leader are shown. **d**, Views of the foreign DNA- (left) and repeat-bound (right) faces of Cas1-2/3 are shown in surface representation and colored by columbic potential. For clarity the highly electronegative DNA is shown in cartoon representation. Labels highlight highly basic and conserved surfaces of each Cas1-2/3 subunit that accommodate the packing of four dsDNA helices in proximity around the Cas2 homodimer (**Extended Data Fig. 3**). IHF heterodimers are shown in 100% transparency for clarity. The phosphate-to-phosphate distances of DNA helices packed around Cas2 are noted.

The structure reveals two prominent DNA bends that protrude at right angles from Cas1-2/3 (**Fig. 1c,d**). An IHF heterodimer is wedged at the apex of each DNA bend, consistent with IHF’s well-defined role in DNA bending ^38^. These two DNA protrussions extend ∼75 Å from the Cas1-2 core. Flexablity of these DNA extentions limits the resolution of the regions to 4-8 Å (**Fig. 1c,d and Supplementary Video 2**). IHF-mediated bending of the IHF_distal_ site positions the flanking IR sequences as symmetrical DNA pillars, which are recognized by equivalent surfaces on opposite sides of the Cas2 homodimer (**Fig. 3 and Extended Data Fig. 3b and 4c**) ^22^. Cas2 binding to these DNA pillars traps foreign DNA on one face of the Cas1-2/3 integrase. Further, Cas2 bends the IRs and steers downstream DNA away from Cas1-2/3, which would project the downstream CRISPR repeat away from the Cas1-2/3 integrase (**Fig. 1d**). However, Cas1-2/3 and IHF cooperate to constrict the DNA around the IHF_proximal_ site, forming a loop that places the CRISPR repeat into the Cas1a* active site (**Fig. 1d and 3**).

## Foreign DNA constrains the Cas2/3 linker against conserved Cas1 surfaces

The type I-F Cas2 and Cas3 subunits are connected by a 20 amino acid disordered linker (residues 90-110) ^4,28,29,55^. The structure explains how foreign DNA constrains the Cas2/3 linker against conserved surfaces of Cas1, which suggests that foreign DNA-binding either initiates, or stabilizes the Cas3 rotation (**Fig. 2a**) ^54,55^. The constrained Cas2/3 linker positions the HD nuclease domain of Cas3 (residues 111-374) against the Cas1-Cas1 interface, and facilitates Cas3 interactions with the IRs (**Fig. 2a and Extended Data Fig. 2 and 4c**). The foreign DNA and amino acids in the Cas2/3 linker contact conserved residues in type I-F Cas1 proteins (**Fig. 2d,e and Extended Data Fig. 2c**). Polar residues in the Cas2/3 linker may assist the binding or splaying of the foreign DNA duplex at the conserved histidine wedge (H25) in Cas1 (**Fig. 2b,c and Extended Data Fig. 3**). Mutation of the histidine wedge (Cas1^H25A^) decreases Cas1-2/3 integration activity of foreign DNA that has either fully complementary or splayed DNA ends (**Extended Data Fig. 5 and 6a,b**). The integration defect on substrates with splayed ends suggest that H25 is more than a simple wedge that pries apart the ends for foreign DNA ^10^. The histidine steers the 3’-ends down a positively charged channel that positions each 3’-hydroxyl into Cas1 active sites on opposite ends of the complex (**Fig. 2 and Fig. 4a,d**), whereas the 5’-ends of the protospacer DNA are directed towards the back face of the Cas3 HD domain (**Fig. 2**) ^9,10,56^.

**Figure 4.**
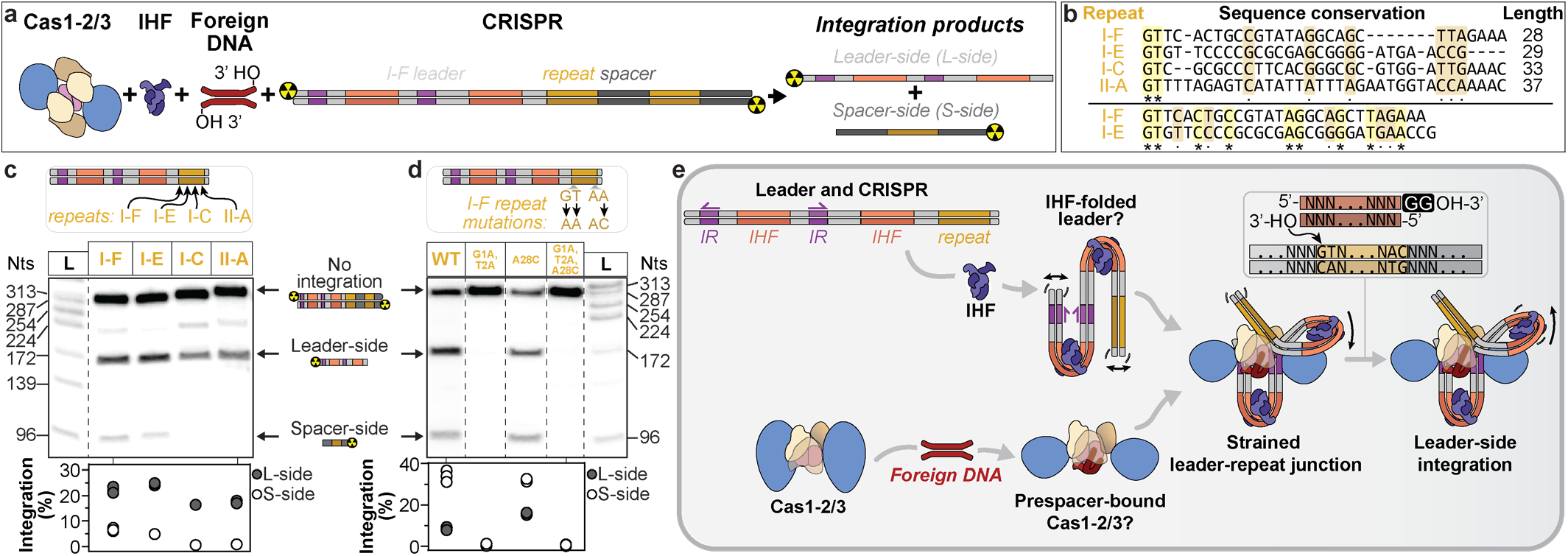
Sequence motifs in the leader and IHF proteins facilitate Cas1-2/3-based integration into diverse repeat sequences. **a**, Scheme of reactants and products of *in vitro* CRISPR integration assays (**Extended Data Fig. 6,7 and 9**). **b**, Four CRISPR repeats used in the integration assays. A gapped sequence alignment highlights two identical (asterisks) and six similar (dots) positions. An un-gapped sequence alignment reveals nine identical nucleotide positions between the I-F and I-E repeats. All four repeats have different internal palindromes and GC-content (**Extended Data Fig. 8b**). **c**, Endpoint integration reactions with CRISPR repeat-swapped mutants, resolved on denaturing polyacrylamide gels. One of three representative gel images is shown (**Extended Data Fig. 7**). Quantification of leader- (grey circles) or spacer-side (white circles) integration events from all three replicate gels (**Extended Data Fig. 7**). The reactions were performed in triplicate, each dot represents one reaction, and some dots overlap. **d**, The 4-minute time point of time-course integration reactions with I-F repeat mutants, resolved on denaturing polyacrylamide gels. One of three representative images is shown (**Extended Data Fig. 9**). Quantification of leader- (grey circles) or spacer-side (white circles) integration events from all three replicate gels (**Extended Data Fig. 9**). **e**, CRISPR integration model. IHF-mediated folding of the genome presents IRs as symmetric DNA pillars that recruit foreign DNA-bound Cas1-2/3. Cas3 domains of Cas1-2/3 must rotate away from Cas2 to expose IR binding sites on Cas2. Cas1-2/3 and IHF cooperate to fold DNA into a loop, docking the leader-repeat junction at the Cas1 active site. Foreign DNA integration at the leader-repeat junction nicks the DNA duplex, releasing tension in the DNA duplex and inhibiting the reverse disintegration reaction (**Extended Data Fig. 4**) ^33,34^. 5’ GT dinucleotides are required for efficient leader- and spacer-side integration, but no strict sequence requirements are necessary in the rest of the repeat.

## Cas2 homodimers recognize and bend inverted repeat sequences

The structure of the type I-F CRISPR integration complex reveals that Cas2 is the homodimer that binds the IRs (**Fig. 3a**). Mutations that scramble the order of nucleotides in either the IR_distal_ or IR_proximal_ motifs limit Cas1-2/3-mediated integration ^22^. While Cas2 doesn’t make extensive sequence-specific contacts with nucleobases of the IR, a single residue (Cas2^R55^) intercalates in the minor groove, and may participate in recognizing two conserved bases in the 10 bp-long motif (**Fig. 3a and Extended Data Fig. 4c,e**). However, there is insufficient density for the R55 sidechain to confidently assign contacts. Other conserved Cas2 residues (i.e., K11, R12, and N56) form additional hydrogen bonds with the phosphate backbone of one DNA strand in each IR (**Fig. 3a and Extended Data Fig. 4c**). Mutation of these Cas2 residues (Cas2^K11D,R12E^, Cas2^R55E,N56D^, Cas2^K11D,R12E,R55E,N56D^) prevents Cas1-2/3-mediated DNA integration (**Extended Data Fig. 5 and 6c,d**). Cas2 acts as a wedge that induces a 25-35° bend in the DNA upstream of IR_distal_ and downstream of IR_proximal_ (**Fig. 3a**). These flared IRs lean against basic residues (K381, R393, K397) on the back surface of Cas3 (**Fig. 3 and Extended Data Fig. 3 and 4c**). In sum, these observations reveal that the IR DNA sequences are primarily recognized by Cas1-2/3 through shape readout rather than base readout ^39^.

## Cas1-2/3 accommodates four dsDNA helices that surround the Cas2 homodimer

Cas2 is a cube-shaped homodimer at the center of the Cas-integrase. The Cas2 cube is flanked by Cas1 homodimers to form an elongated DNA binding platform that interacts with the CRISPR repeat on one face and the foreign DNA on the other (**Fig. 3**). Unique to the type I-F Cas1-2/3 integration complex, the IRs occupy the last two accessible surfaces of the Cas2 cube (**Fig. 3a**). Positively charged surfaces on Cas1-2/3 bind and shield negatively charged DNA, which enables the packing of four DNA helices around the small Cas2 homodimer (**Fig. 3b and Extended Data Fig. 3**). The foreign DNA-binding face of Cas2 has two electronegative pillars of leader DNA that straddle the foreign DNA, such that major grooves of the leader DNA pillars are clamped against major grooves of the foreign DNA. The two DNA pillars continue past Cas2 to flank the Cas1 active sites (**Fig. 3b**). At the IHF_proximal_ loop, Cas3 packs the leader against the Cas1-bound repeat, decreasing the phosphate-to-phosphate distances between these helices to ∼11-12 Å. Although the latter two-thirds of the CRISPR repeat could not be resolved, the repeat’s trajectory suggests it will follow a path that threads between the distal leader DNA duplex and the 3’-hydroxyl of the foreign DNA that rests in the Cas1b* active site (**Fig. 3b**). Collectively, Cas1, the Cas2/3 linker, and Cas3 accommodate four DNA helices (IR_distal_, IR_proximal_, foreign DNA, and repeat) around the central Cas2 homodimer to facilitate site-specific integration.

## IHF and the leader sequence facilitate integration into diverse repeat sequences

The structure suggests that Cas1-2/3 is guided to the first repeat of the CRISPR by IHF-mediated folding of the I-F leader, rather than direct recognition of the repeat sequence (**Fig. 1d and Extended Data Fig. 4b,d**). To determine if or how the repeat sequence impacts integration, we measured the efficiency of Cas1-2/3-catalyzed integration into DNAs containing either a I-F, I-E, I-C or II-A repeat downstream of a I-F leader (**Fig. 4a-c**). The type I-F leader supports Cas1-2/3-catalyzed leader-side integration at repeats derived from I-E, I-C, and II-A CRISPR loci (**Fig. 4a-c and Extended Data Fig. 7 and 8**). Integration efficiency at non-native repeats is not correlated with sequence similarity to the I-F repeat or with GC-content (**Extended Data Fig. 8b**). Instead, integration efficiency is correlated to the length of the repeat. I-F and I-E repeats are similar in length (28 and 29 bps, respectively), whereas the I-C and II-A repeats are 0.5 to 1 full DNA turns longer than the I-F repeat (33 and 37 bp, respectively) (**Fig. 4b and Extended Data Fig. 9a,b**). While leader-side integration is robust with different repeats, spacer-side integration is ∼3.5-fold slower for the I-E repeat, and undetectable for the longer repeats (**Extended Data Fig. 7 and 9a,b)**.

As expected, Cas1-2/3 does not catalyze integration at a I-F repeat downstream of a scrambled I-F leader, nor does Cas1-2/3 does catalyze integration at I-E, I-C or II-A repeats downstream of their respective leaders ^22^ (**Extended Data Fig. 7**). Cas1-2 sequences are diverse, such that leader-interacting residues are only conserved in a subset of proteins within a given CRISPR subtype ^22^. For example, the I-F Cas1 protein lacks residues required to interact with the I-E leader ^22^. Further, the I-E, I-C and I-F leaders have distinct nucleotide spacings between the leader motifs and the repeat. These nucleotide spacings impart a unique shape to the DNA, which is critical for integration.

These experiments were performed with foreign DNA substrates either with or without a PAM (**Extended Data Fig. 7**). Cas1-2/3 integration of PAM-containing DNA is more specific, but the conclusions are otherwise consistent between the two substrates. The PAM must be trimmed by an ancillary nuclease before Cas1-2/3 can catalyze spacer-side integration, therefore we focused our discussion on results from the trimmed foreign DNA to compare differences in spacer-side integration (**Extended Data Fig. 7a-d**) ^14–16^. Collectively, these integration experiments indicate that leader sequences and host factors dictate site-specific integation of foreign DNA at diverse DNA target sites, providing insight into the evolution of new CRISPR systems in nature.

## 5’ GT dinucleotides in CRISPR repeat sequences are critical for Cas1-mediated integration

Repeat sequences are strongly conserved within CRISPR subtypes, but vary in sequence and length between subtypes ^57,58^. However, in the small subset of repeats tested above, we noticed that the 5’ GT is conserved. To determine if conservation of the 5’ GT is a coincidence or a more widely conserved feature of repeats, we performed a bioinformatic analysis consisting of 24,940 CRISPRs. This bioinformatic analysis reveals that a 5’ GT dinucleotide is broadly conserved at the leader-side of the repeat, and conserved in some CRISPR systems at the spacer-side of the repeat (**Extended Data Fig. 4g**). Therefore we hypothesized that the 5’ GT dinucleotide is a base-specific determinant for leader-side integration. To test this hypothesis, we mutated the 5’ GT (G1A, T2A) and repeated the integration assays. The 5’ GT to AA mutation ablates both leader- and spacer-side integration, indicating that the 5’ GT is essential and that leader-side integration is a prerequisite for spacer-side integration (**Fig. 4d**). Since Cas1 requires a 5’ GT at the leader-side of the repeat, we hypothesized that introducing a 5’ GT at the spacer-side of the repeat would increase spacer-side integration efficiency. To test this hypothesis, we replaced adenosine 28 of the I-F repeat with cytosine (A28C) and repeated the integration assays. The A28C mutation increases the rate and amount of spacer-side integration by ∼2-fold, relative to the WT I-F repeat (**Fig. 4d**). We do not detect integration into a I-F repeat that lacks a 5’ GT at the leader-side even if the repeat contains a 5’ GT at the spacer-side. This result further supports our conclusion that leader-side integration is a prerequisite for spacer-side integration (**Fig. 4d**). We examined the structure of the Cas1 active site to determine whether the 5’ G is directly recognized by protein contacts. The Cas1 residue E184 is within 4 Å of the 5’ G (**Extended Data Fig. 4b,d**). However, a Cas1^E184A^ mutation destabilizes the complex, decreasing the amount of Cas1 subunits per Cas1-2/3 complex (**Extended Data Fig. 5**). Therefore, the decrease in integration activity of the Cas1 ^E184A^-2/3 complex cannot be solely attributed to a decrease in recognition of the repeat (**Extended Data Fig. 9d,f**). Collectively, these data reveal that the 5’ GT is a conserved feature necessary for integration in most CRISPR systems, though no available structure provides a mechanism for direct recognition of the 5’ GT of the repeat ^5,11,15,59^.

## Conclusions

Here we demonstrate that Cas1-2/3 and IHF fold DNA into a structure that is necessary for site-specific integration of foreign DNA into CRISPRs. IHF proteins are highly expressed and most IHF binding sites are thought to be occupied *in vivo* ^60^. Therefore, IHF may pre-fold the CRISPR leader into a "landing pad" that recruits foreign DNA-bound Cas1-2/3 (**Fig. 4e**) ^22,38^. The Cas3 and Cas1 domains of the Cas1-2/3 complex are arranged like petals of a closed flower around the central Cas2 homodimer, such that the Cas3 domains occlude two of the four DNA binding surfaces on Cas2, which precludes interactions with the leader (**Fig. 1b,d**). Foreign DNA binding to Cas1-2/3 physically constrains the Cas2/3 linker against the Cas1 homodimer, pulling the Cas3 HD domain against the Cas1-Cas1 interface (**Fig. 2, Extended Data Fig. 2 and Supplementary Video 1**). The 100-degree rotation of each Cas3 simulates the motion of a bloomed flower and exposes DNA binding sites on Cas2 that interact with each of the inverted repeat (IR) motifs in the leader (**Fig. 1, 3, Extended Data Fig. 2,3**). Cas1-2/3 and IHF proteins fold DNA around the leader-repeat junction into a 260° loop that docks the first CRISPR repeat into the Cas1 active site under tension (**Fig. 1, 4, Extended Data Fig. 3, 4**). The structure suggests that Cas1-mediated strand-transfer releases tension in this DNA loop, which may prevent disintegration of an otherwise isoenergetic strand-transfer reaction, and thereby favour complete integration (**Extended Data Fig. 4f**). A similar mechanism has been proposed to favour complete integration in other systems, where both strand transfer events occur simultaneously ^61,62^.

We show that Cas1-2/3, IHF and the I-F leader facilitate leader-side integration at four different repeat sequences (**Fig. 4b-c**). These repeats are diverse in sequence identity, length, palindrome and GC-content, but they share a 5’ GT (**Fig. 4b and Extended Data Fig. 8b**). To determine whether the 5’ GT is a universal feature of CRISPR repeats we analyzed 24,940 CRISPRs. This analysis reveals that CRISPR repeats contain a strongly conserved 5’ GT at the leader-end, and that a 5’ GT is also conserved at the spacer-end of repeats from several CRISPR systems (**Extended Data Fig. 4g**). We demonstrate the 5’ GT is critical for leader-side integration and that introducing a 5’ GT at the spacer-end of the repeat increases spacer-side integration. The broad conservation of 5’ GT is consistent with previous reports that type I-A, II-A, and I-E systems require a 5’ G for integration ^11,63^. Collectively, these data suggest that Cas1 proteins retain a shared sequence preference for a 5’ G or a 5’ GT, and lack strict sequence requirements for the central body of the repeat (**Fig. 4b,c**) ^64^. The lack of strict sequence requirements may be advantageous because the CRISPR repeat is at the nexus of foreign DNA integration, processing of the transcribed CRISPR, and loading the mature crRNA into the surveillance complexes (e.g., Cascade, Cas9).

Genetic parasites commonly escape CRISPR-based immunity through point mutations ^65^. To counter escape mutants, many CRISPR-Cas systems use existing spacer sequences to enhance the acquisition of new spacers from the same foreign genetic element via “primed” acquisition ^4,30–37^. In some examples of primed acquisition, the Cas3 nuclease/helicase degrades CRISPR-targeted DNAs into ssDNA fragments enriched in PAM-containing termini ^66^. Cas1-2 has been proposed to anneal complementary ssDNA fragments and integrate these into CRISPRs ^14,30,31,66^. Single-molecule co-localization and bulk immunoprecipitation suggest that the type I-E Cas1-2 integrase is recruited to a Cas3-Cascade-target DNA complex to facilitate primed acquisition ^34,67^. The structure of the type I-F integration complex reveals conformational changes in Cas3 that may enable interactions with DNA-bound Cascade (**Fig. 5a**) ^36,68^. Cascade improves integration efficiency and fidelity *in vivo* ^68,69^, and the structure suggests a model for the formation of a primed acquisition complex (Cas1-2/3-Cascade-target DNA) that transfers new foreign DNA fragments to the integrase. Additional structures will be necessary to clarify the mechanism(s) of primed adaptation.

**Figure 5.**
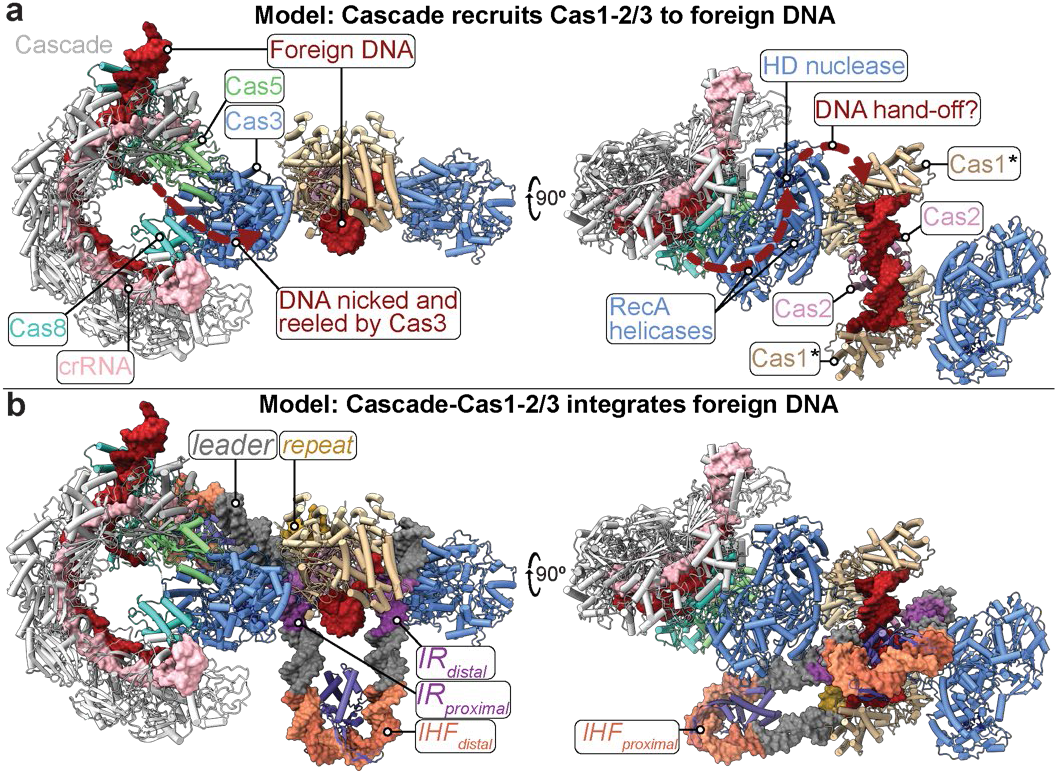
I-F CRISPR integration complex suggests a mechanism for primed acquisition and Cascade impact on integration. **a**, Cascade (grey) bound to foreign DNA (red) displays a Cas8 helical bundle (turquoise) that recruits Cas2/3. Cascade-recruited Cas3 degrades foreign DNA into fragments that are captured by the Cas1-2 integrase for subsequent integration ^37,38,45,46,79–83^. The Cas1-2/3 integration complex can be docked onto dsDNA-bound Cascade with minimal clashing and suggests a model for the formation of a primed acquisition complex that facilitates rapid adaptation to genetic parasite variants 41,47,81. **b**, Recruitment of the Cascade-Cas1-2/3 complex to IHF-folded CRISPR leader DNA is not predicted to form any new clashes, suggesting a mechanism for the role of Cascade in facilitating integration *in vivo* 72,73. Note that a total of two Cascade complexes can be docked onto the Cas1-2/3 integration complex (one per each Cas3) without introducing additional clashing, a single is shown above for clarity.

A comparison of the I-E and I-F integration complexes with a structure of the λ-phage excision complex, reveals structural similarities and differences. In I-E systems, the IHF protein folds the leader DNA to present an upstream motif to a lobe of Cas1. In contrast, the I-F structure highlights extensive cooperation between IHF and Cas1-2/3 in bending the leader into an energetically strained conformation that may increase the specificity of CRISPR recognition (**Fig. 6a,b**). This cooperation includes Cas1-2/3 kinking the IR DNA motifs presented as parrallel DNA pillars by IHF, and Cas1-2/3 constricting the IHF_proximal_ 180° bend ito a 260° (**Fig. 6b**). Further, the sequestration of the IR-binding surface of Cas2 suggests a unique structural mechanism that prevents Cas1-2/3 interactions with the leader until foreign DNA binding induces a rotation of Cas3 (**Fig. 4e, Supplementary Video 1 and 3**) ^54^.

**Figure 6.**
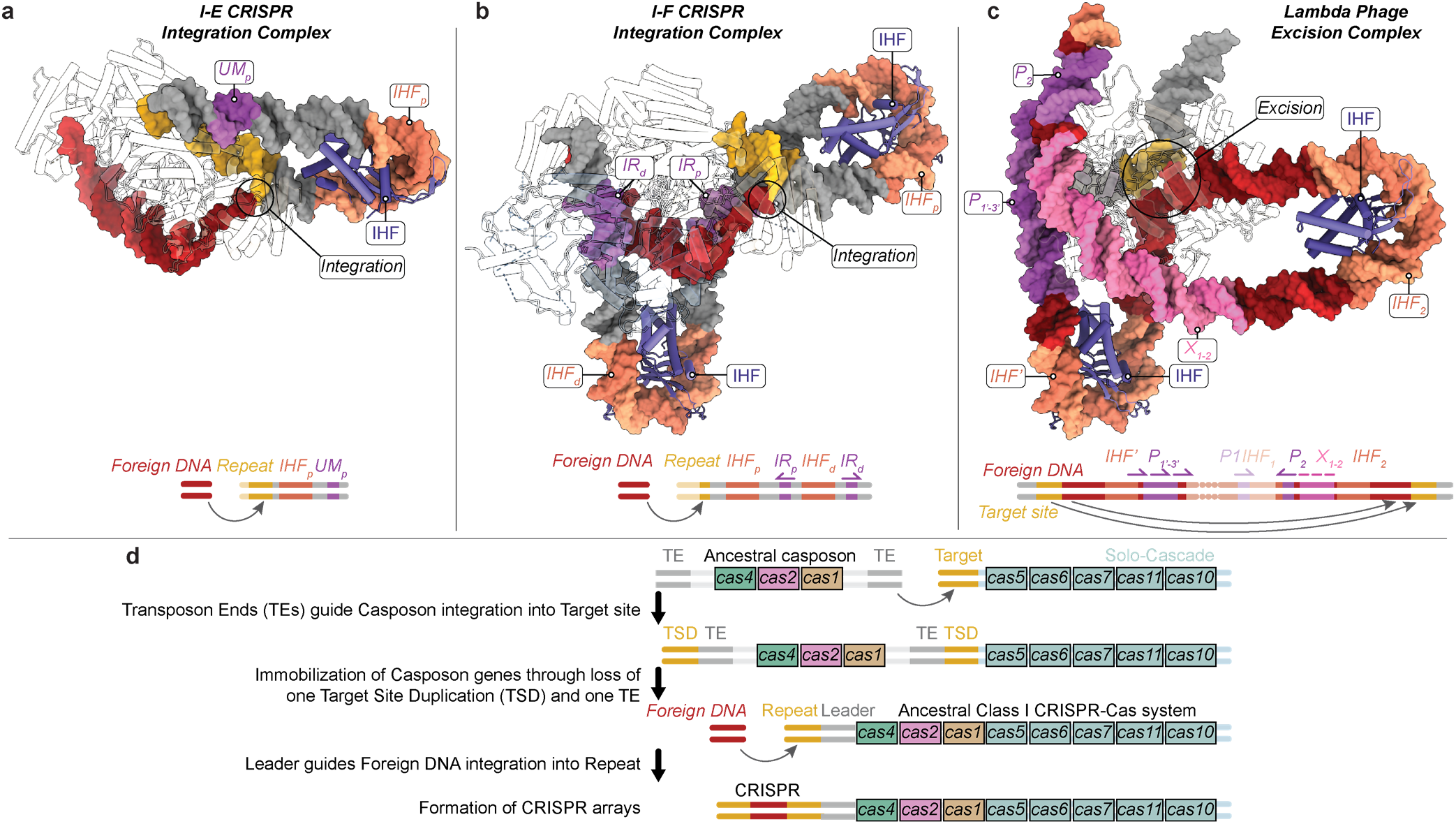
DNA is a flexible scaffold that controls DNA mobilization. Structures for the I-E CRISPR integration complex **(a**), I-F CRISPR integration complex (**b**), and Lambda phage excision complex (**c**). DNA shown as a surface, IHF (purple) and all other proteins shown as transparent cartoons. Integration and excision sites, along with DNA motifs that regulate DNA mobilization, are labeled and colored according to the schematic (bottom). **d**, A revised evolutionary scenario for CRISPR-Cas systems arising from a transposon that uses Cas1 as an integrase (“Casposon”) is proposed ^51^. An ancestral Casposon encoding Cas1, Cas4 and Cas2 is shown. Transposon ends with DNA motifs that regulate transposition (grey) facilitate Casposon integration into a DNA target site near an operon encoding a generic type III Cascade complex (light green). Loss of one transposon end and target site duplication immobilizes the Casposon genes. The remaining transposon end (Leader) was repurposed for foreign DNA integration into the target site (repeat), forming CRISPR arrays with a repeat-spacer-repeat architecture.

Many CRISPR leaders contain multiple IHF binding sites and subtype-specific motifs that are reminiscent of motifs found at the ends of transposons, phages, and plasmids, which facilitate integration, excision, or recombination complex assembly (**Extended Data Fig. 8a**) ^3–7,22,40–53^. For example, the left and right ends of the λ-phage genome contain three IHF binding sites, five copies of a second DNA motif (P1-2,1’-3’), and three copies of third DNA motif (X1,1.5,2), that recruit λ-phage (Int, Xis) and non-λ-phage (IHF, Fis) proteins (**Fig. 6c**)^7^. These proteins use DNA as a flexible scaffold that organizes enzyme active sites and DNA substrates to facilitate either the integration, or excision, of λ-phage DNA from the bacterial genome (**Fig. 6c**). Similarly, the I-F CRISPR leader motifs recruit Cas (Cas1-2/3) and non-Cas (IHF) proteins, that bend DNA into a flexible scaffold to organize enzyme active sites and DNA substrates in space, facilitating the integration of foreign DNA into the bacterial genome (**Fig. 6b**). Diverse systems thus use DNA as a flexible scaffold to regulate the isoenergetic mobilization of DNA (**Fig. 6a-c**).”

The CRISPR adaptive immune system has been proposed to have originated through the domestication of a transposon that used a Cas1 homolog to catalyze integration (Casposon) ^1^. The pathway for the domestication of proteins from different operons to form the Cas components of CRISPR-Cas system has been previously proposed ^1,70^. However, the domestication or evolution of DNA elements that gave rise to the CRISPR repeat and leader has remained unclear ^1,70^. The similarities in motif architectures between CRISPR leaders and transposon DNA ends suggests that the Capsoson DNA end may have been domesticated to form the primordial CRISPR “leader” (**Fig. 6d**). Similarly, the target DNA site that the ancient Casposase integrated at may have been co-opted to form the first CRISPR “repeat” (**Fig. 6d**). Originally, the transposon end directed Casposase to integrate Casposon (foreign) DNA into the target site. Now, the domesticated leader directs Cas1-2 to integrate foreign DNA into the domesticated repeat (**Fig. 6d**).

Diverse DNA mobilizing enzymes across the tree of life co-opt DNA folding to regulate DNA mobilization ^3–7^. In sum, these data provide a mechanistic understanding for the role of DNA as a flexible scaffold that controls DNA mobilization. These insights are critical to developing applications of DNA-mobilizing enzymes in gene therapy, genetic engineering, and chronological DNA recordings ^6,53,71–75^.

## Methods

### Nucleic acid preparation

Four single-stranded DNAs (**Supplementary Table S1**) were synthesized (IDT) and resuspended in 1x TE buffer (10 mM Tris-HCl pH 8, 1 mM EDTA) before being used to assemble the structure of the type I-F integration complex, the assembly is detailed in a below section. The splayed foreign DNAs used in integration assays (**Supplementary Table S1**) were synthesized and resuspended in 1x TE buffer before use. To make ^32^P-labeled CRISPR integration substrates, the sequences consisting of the leader and CRISPR arrays were first synthesized and cloned into pUC57 (Genscript). These plasmids have been made available on Addgene (**Supplementary Table S1**). These plasmids were transformed into chemically competent *E. coli* DH5α cells and the transformed cells were plated onto LB agar plates containing 100 µg/mL Ampicillin. These cells were cultured in LB media and plasmids were purified using ZymoPURE II Plasmid Midiprep kit (Zymo Research). Each plasmid was then digested with EcoRI-HF and BamHI-HF (NEB) restriction enzymes, and the 294-383 bp inserts of interest were separated from the vector backbone by agarose gel electrophoresis. The gel segments containing the DNA inserts of interest were excised and DNA was purified using a Zymoclean Gel DNA Recovery kit (Zymo Research) (**Supplementary Table S1**). The 5’ ends of the CRISPR leader and array fragments were dephosphorylated using Quick Calf Intestinal alkaline Phosphatase (NEB), and the DNAs were purified away from protein using a DNA Clean & Concentrator kit (Zymo Research). Both 5’ ends of 1 pmole of the CRISPR leader and array fragments were then labelled with ^32^P, by incubation with 4 pmoles of [γ-^32^P]ATP (PerkinElmer) by polynucleotide kinase (NEB) in 1x PNK buffer at 37°C for 45 minutes. PNK was heat-denatured by incubation at 65°C for 20 minutes. Spin column purification (G-25, GE Healthcare) was used to remove unincorporated radioactive nucleotides and to buffer exchange DNAs into 1x TE buffer.

### Cas1 & Cas2/3 Mutagenesis

The plasmid used to express Cas1 and Cas2/3 (Addgene #89240) was PCR amplified with mutagenic primer pairs using Q5 polymerase (NEB) (**Supplementary Table S1**). The parental template plasmid was digested with DpnI (NEB) and the PCR amplified products were purified using the DNA Clean and Concentrate kit (Zymo). The purified DNA was 5’ phosphorylated with T4 Polynucleotide Kinase (NEB) and ligated with homemade T4 DNA ligase. The ligation reactions were transformed into DH5α cells. The Cas2 K11D, R12E, R55E, N56D mutant was prepared using the Cas2 K11D, R12E mutant as the template with the PCR primers for Cas2 R55E, N56D in the mutagenic PCR reaction. Plasmids for expressing the mutant Cas1-2/3 complexes, Cas1H25A-2/3, Cas1E184A-2/3, Cas1-2K11D,R12E/3, Cas1-2R55E,N56D/3, and Cas1-2K11D,R12E,R55E,N56D/3 have been deposited at Addgene (#200213, #200214, #200216, #200217, #200218).

### Protein purification

*P. aeruginosa* IHF heterodimer was purified as previously described ^22^. Briefly, 6xHis-tagged IHFα and StrepII-tagged IHFβ were co-expressed in *E. coli* BL21(DE3). Cell pellets were lysed by sonication in IHF lysis buffer (25 mM HEPES-NaOH pH 7.5, 500 mM NaCl, 10 mM Imidazole, 1 mM TCEP, 5% Glycerol), supplemented with 0.3x Halt Protease Inhibitor Cocktail (ThermoFisher), at 4°C. Lysate was clarified by two rounds of centrifugation at 12,000 rpm for 15 minutes, at 4°C. His-tagged IHF was captured on HisTrap HP resin (Cytiva), and eluted with 500 mM Imidazole. Affinity tags were cleaved using PreScision protease, and the PreScision protease and remaining 6x-His-IHFα were removed by affinity chromatography using HisTrap HP resin (Cytiva). Untagged IHF heterodimer was then further purified on Heparin Sepharose (Cytiva) and eluted with a linear gradient to a buffer containing 2 M NaCl. Fractions containing IHF heterodimer were concentrated and further purified by SEC (<u>S</u>ize <u>E</u>xclusion <u>C</u>hromatography) on a Superdex 75 column (Cytiva) equilibrated in IHF buffer (25 mM HEPES-NaOH pH 7.5, 200 mM NaCl, 5 % Glycerol) (**Extended Data Fig. 1a**). IHF overexpression plasmids have been previously described and deposited at Addgene (#149384, #149385) ^22^.

*P. aeruginosa* Cas1-2/3 heterohexamer complexes (wildtype and mutants) were purified as previously described ^22^. Briefly, StrepII-tagged Cas1-Cas2/3 was overexpressed in *E. coli* BL21(DE3). Cell pellets were lysed via sonication in Cas1-2/3 lysis buffer (50 mM HEPES pH 7.5, 500 mM KCl, 10% Glycerol, 1 mM DTT), supplemented with 0.3x Halt Protease Inhibitor Cocktail (ThermoFisher), at 4°C. Lysate was clarifed as above. StrepII-tagged Cas1-Cas2/3 complexes were affinity purified on StrepTrap HP resin (GE Healthcare) and eluted with Cas1-2/3 Lysis Buffer containing 3 mM desthiobiotin (Sigma-Aldrich). Eluate was concentrated at 4°C (Corning Spin-X concentrators), before purification over a Superdex 200 size-exclusion column (Cytiva) equilibrated in 10 mM HEPES pH 7.5, 500 mM Potassium Glutamate, and 10% Glycerol (**Extended Data Fig. 1c and 5e**).

### *In vitro* integration assays

End-point integration reactions were performed in triplicate using 300 nM of splayed foreign DNA fragments, containing or lacking a PAM (IDT), 200 nM of Cas1-2/3, 300 nM of IHF heterodimer and roughly 1 nM of a given ^32^P-labelled CRISPR variant fragment in Integration Buffer (20 mM HEPES pH 7.5, 150 mM potassium glutamate, 5 mM MnCl_2_, 1 mM TCEP, 1% Glycerol) (**Supplementary Table S1)**. Reactions were assembled on ice and then incubated at 35°C for 20 minutes.Timecourse integration reactions were performed in triplicate using 300 nM of splayed, or fully complementary, foreign DNA fragments lacking a PAM (IDT), 200 nM of Cas1-2/3, 300 nM of IHF heterodimer and roughly 1 nM of a given ^32^P-labelled CRISPR variant fragment in modified Integration Buffer (20 mM HEPES pH 7.5, 150 mM potassium glutamate, 5 mM MnCl_2_, 10 mM TCEP, 1% Glycerol) (**Supplementary Table S1)**. We noticed that IHF and Cas1-2/3 protein stocks exhibited a propensity to precipitate when diluted into pre-chilled buffer to form working dilution stocks. Therefore all working dilutions of IHF and Cas1-2/3 prepared for timecourse assays were made by first diluting the proteins into room-temperature buffer, mixed, and then chilled on ice. Reactions were assembled on ice and then incubated at 20 minutes. Timepoints were taken at 0, 1, 2, 4 and 8 minutes. Reactions were stopped by the addition of phenol. The aqueous (nucleic acid-containing) layer was mixed 1:1 with 2x formamide loading buffer (95% formamide, 20 mM EDTA, 0.05% bromophenol blue, 0.05% xylene cyanol) and then denatured at 95°C for 5 minutes, before resolving the ^32^P-labeled CRISPR substrates and integration products on a 7% (w/v) (29:1 mono:bis) polyacrylamide Urea gel in 1x TBE (100 mM Tris-Borate pH 8.3, 2 mM EDTA). Gels were dried and quantified using a Typhoon phosphorimager (GE Healthcare). The intensity of full-length CRISPR variant, leader-side integration fragments and spacer-side integration fragments were quantified with Multi Gauge v3 (Fujifilm). These readings were then used to calculate leader- and spacer-side integration events as percentages of all events. Images of all gels that resolved integration reactions are shown **Extended Data Fig. 6,7 and 9**), and additional control gels show that Cas1-2/3 is required for integration, and show how the custom ^32^P-labelled ladder was generated by restriction enzyme digestion of ^32^P-labelled CRISPR variant DNAs (**Extended Data Fig. 8**). Timecourse integration data was fit to a plateau followed by one phase association (GraphPad Prism).

### Assembly and purification of I-F integration complex

A total of four single-stranded DNAs (ssDNAs) synthesized to mimic a half-site integration intermediate were annealed in a step-wise manner. 2 nanomoles of ssDNAs mostly corresponding to the sense and anti-sense strands of the CRISPR leader ("strand_1" and "strand_2" were denatured at 100°C and then slow annealed using a PCR program that cooled the samples to 25°C in 6°C over an hour, in 100 µL of hybridization buffer (20 mM Tris-HCl pH 7.5, 100 mM monopotassium glutamate, 5 mM EDTA, 1 mM TCEP). 2 nanomoles of ssDNAs mostly corresponding to the corresponding to the sense and anti-sense of the strands of the foreign DNA ("strand_3" and "strand_4") were slow annealed using the same protocol (**Extended Data Fig. 1a and Supplementary Table S1**). The two sets of annealed DNAs (tube 1: "Strand_1" and "Strand_2"; tube 2: "Strand_3" and "Strand_4") were mixed together, heated to 80°C and then slow annealed using a PCR program that cooled the samples to 25°C in 6°C over an hour, to anneal the complementary sense and anti-sense regions of the CRISPR repeat included in Strand_2 and Strand_3 together. Next, 6 nanomoles of IHF heterodimer in 50 µL of hybridization buffer was warmed to 25°C and then mixed and incubated with the annealed DNAs at 25°C for 10 minutes. Next, 3 nanomoles of Cas1-2/3 in 250 µL of hybridization buffer was warmed to 25°C and mixed with the prepared DNA and IHF mixture, and incubated at 25°C for 10 minutes. The total concentration of monopotassium glutamate in the mixture at this stage was ∼200 mM, due to carryover from the stored protein stocks. This sample was centrifuged at 22,000 g, 4°C for 20 minutes to remove precipitates. The type I-F CRISPR integration complex was then purified on a Superdex 200 10/300 column (Cytiva) equilibrated in SEC buffer (20 mM Tris-HCl pH 7.5, 200 mM monopotassium glutamate, 5 mM EDTA, 1 mM TCEP, 2 % Glycerol). 0.5 ml fractions were individually concentrated and stored. The sixth SEC fraction contained all DNAs and proteins of interest and was further analyzed by cryo-EM (**Extended Data Fig. 1d-f**).

### Cryo-EM sample preparation and data acquisition

Purified integration complex was diluted to a concentration of 1 µM in SEC buffer lacking glycerol (20 mM Tris-HCl pH 7.5, 200 mM monopotassium glutamate, 5 mM EDTA, 1 mM TCEP), such that the final glycerol concentration was 0.2% within 1 hour of freezing. Sample was applied to Quantifoil R2/2 Cu 200 mesh grids that were glow discharged using 15 mA for 15 seconds with a 10 second hold (easiGlow, Pelco). 4 µl of diluted integration complex was applied to the grids, and then the grids were blotted for 5-6 seconds using Vitrobot^TM^ Filter paper (Electron Microscopy Sciences) with a blot force of 6, at 100% humidity, 8°C, followed by plunge freezing into liquid ethane using a Vitrobot (Mk. IV, ThermoFisher Scientific). A preliminary dataset of 230 movies was collected on Montana State University’s Talos Arctica transmission electron microscope (ThermoFisher Scientific), with a field emission gun operating at an accelerating voltage of 200 kV using parallel illumination conditions ^76^. Movies were acquired using a Gatan K3 direct electron detector, operated in electron counting mode applying a total electron exposure of 50 e-/Å^2^ over 50 frames (3.995 s exposure, 0.08 s frame time). The SerialEM data collection software was used to collect micrographs at 36,000-fold nominal magnification (1.152 Å/pixel at the specimen level) with a nominal defocus set to 0.5 µm - 2.0 µm ^77^. Stage movement was used to target the center of four 2.0 µm holes for focusing, and image shift was used to acquire high magnification images in the center of each of the holes. An preliminary reconstruction was determined from a curated set of 160 images that had CTF fits less than 9 Å and a full-frame motion less than 40 pixels. Briefly, a round of blob picking (150-270 Å) followed by 2D classification was used to identify 2D classes used as templates for template picking in cryoSPARC ^78^. Template picking identified 33,403 initial particles from the above 160 images. 2D classification of these 33,403 particles into 50 classes was used to identify 5 classes with strong structural features containing 4,002 particles. Non-uniform refinement of these 4,002 particles resulted in a ∼14.7Å reconstruction that appeared to contain a complete integration complex, therefore new grids were prepared as described above and shipped to National Center for CryoEM Access and Training (NCCAT) and the Simons Electron Microscopy Center located at the New York Structural Biology Center (NYSBC) for additional data collection (**Table 1**). At NCCAT, grids were imaged using a 300 kV Titan Krios G3i (Thermo Fisher Scientific) equipped with a GIF BioQuantum and K3 camera (Gatan). 10,740 images were recorded with Leginon ^79^ (Suloway et al., 2005) with a calibrated pixel size of 0.5335 Å/px (micrograph dimension of 11520 x 8184 px) over a nominal defocus range of −0.7 μm to −2.1 μm and 20 eV slit. Movies were recorded in “super-resolution mode” (native K3 camera binning 1) with subframes of 50 ms over a 2.5 s exposure (50 frames) to give a total exposure of ∼69 e-/Å^2^ (**Table 1**).

### Cryo-EM image processing

Patch motion correction and patch CTF correction were performed in cryoSPARC ^78^. 3,792 of 10,740 total images (*CTF < 8Å, Full-frame motion < 30Å*) were processed first to build an initial template. Blob picking was used to pick particles with diameters ranging from 120-280 Å. These ∼1.8 million particles were extracted, Fourier-binned 2×2, and then subjected to 2D classification (*Custom parameters: Initial classification uncertainty factor = 3; Number of online-EM iterations = 30; Batchsize per class = 200*) (**Extended Data Fig. 1g**). Particles from 82 of the 200 2D classes were selected for an initial round of *ab initio* reconstruction and heterogeneous refinement. Particles from 1 of 5 of these classes were selected for a second round of *ab initio* reconstruction and heterogeneous refinement. Particles from 1 of 3 of these classes (174,000 particles) were selected for Non-uniform refinement (*Custom parameters: Optimize per-particle defocus = true; Optimize per-group CTF params = true*) to create an initial reconstruction with a resolution of ∼3.7Å, that was used to calculate templates ^80^. These templates were used to choose particles from 9,858 of 10,740 total images (CTF < 8Å cutoff). These ∼5.85 million particles were classified into a total of 6 classes by Heterogeneous refinement, that were seeded with 1 good volumes and 5 junk volumes taken from the above Heterogeneous refinement analyses. The ∼1.31 million particles in the single selected class, were passed through a round of 2D classification (*Custom parameters: Batchsize per class = 200*) (**Extended Data Fig. 1h**). ∼1.29 million particles from 49 of the 50 2D classes were selected for a round of *ab initio* reconstruction followed by heterogeneous refinement into two classes. ∼1.1 million particles from one of these two classes were subjected to 3D classification into 4 classes (*Custom parameters: Batchsize per class = 20,000; Initialization mode = PCA; Target resolution = 2Å; Particles per reconstruction = 500; Class similarity = 0.3*), followed by separate Non-uniform refinements of particles from each of these four classes (*Custom parameters: Optimize per-particle defocus = true; Optimize per-group CTF params = true*)^80^. The final set of 366,794 particles were re-extracted and re-centered, Fourier-binned 2×2, and subjected to Non-uniform refinement to generate a final reconstruction refined to a global resolution of 3.48 Å based on the 0.143 FSC cutoff (**Extended Data Fig. 1h-k and Table 1**) ^81^. The 3D FSC was calculated using webserver (3dfsc.salk.edu)^82^.

### Model building and validation

The map was sharpened from two half-maps using the local anisotropic sharpening job in Phenix ^83^. The published structure of the *P. aeruginosa* Cas1 homodimer was used as starting models^56^, because Colabfold consistently failed to predict the alternative fold that one Cas1 subunit adopts to form the assymetric homodimer interface ^84^, even when provided template structures. Whereas the Colabfold-predicted models for the *P. aeruginosa* IHF heterodimer, and the *P. aeruginosa* Cas2/3 subunit were used as starting models. The conformation of the DNA sequences within the *E. coli* IHF-DNA co-crystal structure (PDB: 1IHF) ^38^ was used as starting models for DNA segments within the IHF_distal_ and IHF_proximal_ DNA bends. For all other double stranded DNA segments, B-form DNA was used as a starting model. Single-stranded DNA segments were built in *de novo*. Protein and DNA segments were individually rigid-body fitted into the EM density map. The relative orientation of the Cas2, Cas2/3 linker and Cas3 domains were corrected by real-space refinement into the EM density map in WinCoot ^85^. The ReadySet job in Phenix was used to generate hydrogens on all proteins and nucleic acids and prepare the model for further refinement. Then protein and DNA segments were real-space refined in WinCoot ^85^, restrained to ideal geometry, secondary structure and German McClure distance restraints generated in ProSMART from the input models ^86^. The models were iteratively real-space refined in WinCoot and in Phenix using Ramachandran and secondary structure restraints ^83,85^. The starting model was used as a reference model, and harmonic restraints on the starting coordinates were enabled. MolProbity ^87^ and the PDB validation service server (https://validate-rcsb-1.wwpdb.org/) were used to identify problem regions subsequently corrected in WinCoot ^85^. For regions of the reconstruction where side chains are not visible (resolution >4.0Å) the atomic model was truncated to the peptide backbone. For regions of the reconstruction where the backbone was ambigous the sections of the peptide or DNA model were removed. Contacts and hydrogen bonds between residues were identified by ChimeraX v1.4 using the “contacts” and “hbonds” commands respectively, with default parameters ^88,89^. The DNAproDB webserver (https://dnaprodb.usc.edu/) was further used to analyze DNA-protein contacts (**Extended Data Fig. d,e**) ^90^. Structure-guided mutagenesis was used to further validate key Cas1-2/3-DNA contacts in the above biochemical assays.

### Cas1, Cas2/3 and repeat conservation analysis

To build a list of type I-F Cas1 sequences, CRISPRDetect v2.4 with default parameters was used to identify CRISPR arrays within a total of 18,225 bacterial and 376 archaeal complete genomes accessed from the NCBI Assembly database on June 10^th^ of 2019 as previously described ^22,91^. 15,274 high-confidence CRISPR arrays were classified with a CRISPR subtype by CRISPRDetect v2.4 (by matching to a list of repeats with known subtype annotations), and by genetic proximity to subtype-specific *cas* genes (within 20,000 bp). To identify *cas* genes, the 20,000 bp flanking the CRISPR were submitted to PRODIGAL v2.6.3 (default parameters) to predict all potential <u>o</u>pen reading frames (ORFs) ^92^. This ORF database was then used as input to search for *cas* gene clusters with MacsyFinder v1.0.5 ^93^. The following parameters were used: “*macsyfinder --sequence-db <peptide_database> --db-type gembase -d <CRISPR_subtype_definitions> -p <HMM_profiles> -w 50 -vv all*”. HMM profiles and classification definitions used in MacsyFinder were acquired from the local version of CRISPRCasFinder v4.2.20 ^94^. Next, the first repeat and 200 nucleotides upstream of CRISPR arrays (leader) which were classified as Type I-F (1,683 arrays) were collected. A non-redundant list of I-F CRISPR leaders (536 leaders) was generated using CD-HIT v4.8.1 with a 95% identity cutoff ^95^. A local copy of FIMO was used to find significant matches to the position weight matrix representing I-F IHF binging site as previously described ^22,96^. I-F CRISPR arrays that possess more than one IHF site (IHF_proximal_ and/or IHF_distal_) in the leader sequences were extracted for downstream analyses. Cas1 homologs were identified within the 20,000 base-pair flanking regions of extracted 444 I-F CRISPR arrays by using PRODIGAL and MacsyFinder with the same parameters described above. 371 Cas1 homologs associated with Type I-F CRISPRs and possessing at least one IHF site in the leader sequences were identified. A non-redundant list of Cas1 sequences was generated CD-HIT v4.8.1 with a 95% identity cutoff, resulting in 222 sequences ^95^. Sequences smaller than 200 residues and larger than 500 residues were removed, and the remaining 205 sequences were further curated with MaxAlign, which selected a list of 144 unique type I-F Cas1 sequences ^97^. The *P. aeruginosa* PA14 Cas1 sequence was then added to a final list of 145 type I-F Cas1 sequences. To build a list of type I-F Cas2/3 sequences, the *P. aeruginosa* PA14 Cas2/3 sequence was used as an input for HHMER for a search for homologs using 3 iterations, an E-value cutoff of 0.0001, against the UNIREF-90 database ^98,99^. A list of 500 representative sequences was further curated with MaxAlign, to generate a final list of 458 unique Cas2/3 sequences. Type I-F Cas1 and Cas2/3 sequences were aligned using the MAFFT webserver with the E-INS-I iterative refinment methods to result in alignments with the highest number of gap-free sites ^100^.

To build an updated list of CRISPR repeat sequences, CRISPRDetect v3.0 with default parameters was used to identify CRISPR arrays within a total of 25,502 bacterial and 398 archaeal complete genomes and chromosomes accessed from the NCBI RefSeq Assembly database (accessed on June 10^th^, 2021)^91^. This search identified CRISPR loci within 58,864 genomic and plasmid sequences, resulting in 24,940 high-confidence CRISPR loci predictions (array quality score >3). Similar to above, CRISPRDetect annotated the subtype of 14,446 of these CRISPR loci, based on the sequence similarity of the repeats in these loci to known CRISPR repeats. The subtypes of the remaining 10,494 CRISPR loci were determined by their proximity to subtype-specific *cas* genes as described above. 5,321 of the 10,494 unclassified CRISPR loci were assigned a subtype using this protocol, such that 5,173 CRISPR loci remained unclassified. The consensus repeat for each of the 24,940 CRISPR loci, as reported by CRISPRDetect, were used for downstream analyses. To ensure the repeats were arranged in the correct orientation, the 24,940 repeats were grouped by subtype, and each group was individually aligned by MAFFT using the “*--adjustdirection*” parameter. Sequence logos of the first and last three bps of CRISPR repeats were made using Weblogo v3.7.1 for CRISPR subtypes and across all subtypes^101,102^ (**Extended Data Fig. 4g**).

## Supporting information

Supplementary Video 1: Cas1-2/3 undergoes a large conformational change to unveil Cas2-leader binding sites.

Supplementary Video 2: Overview of how the I-F CRISPR leader and IHF guide Cas1-2/3-mediated integration of foreign DNA at the first CRISPR repeat.

Supplementary Video 3: The strained IHF-mediated DNA bends sway in relation to the Cas1-2/3 integrase.

## Reporting summary

Further information on research design is available in the Nature Research Reporting Summary linked to this paper.

## Data availability

The data that support the findings of this study are available from the corresponding author Blake Wiedenheft upon request. Cryo-EM maps were deposited in the Electron Microscopy Data Bank under accession number EMD-29280. The atomic model of the type I-F integration complex was deposited in the PDB under accession number 8FLJ. Plasmids generated in this study are available from Addgene.

## Code availability

Code will be made available upon request and without restriction.

## Acknowledgements

Thanks to members of the B.W. laboratory for feedback and discussions. We thank Dr. Mariusz Matyszewski and Dr. Jeliazko Jeliazkov for helpful discussions. Thanks to Coltran Hophan-Nichols for computational support. A.S-F. is a postdoctoral fellow of the Life Science Research Foundation that is supported by the Simons Foundation. A.S-F. is supported by the Postdoctoral Enrichment Program Award from the Burroughs Wellcome Fund. This work was supported by National Institutes of Health, United States grant 1K99GM147842 (A.S-F.). L.T., A.B.G. is supported by Montana State University’s Undergraduate Scholars Program, and by the NIH NIGMS IDeA program (P20GM103474). This work was performed using the cryo-EM Facility at Montana State University (NSF 1828765 and the M.J. Murdock Charitable Trust). Microscopy was also performed at the National Center for CryoEM Access and Training (NCCAT) and the Simons Electron Microscopy Center located at the New York Structural Biology Center, supported by the NIH Common Fund Transformative High Resolution Cryo-Electron Microscopy program (U24 GM129539), and by grants from the Simons Foundation (SF349247) and NY State Assembly. Research in the Wiedenheft lab is supported by the NIH (R35GM134867), the M.J. Murdock Charitable Trust, a young investigator award from Amgen, and the Montana State University Agricultural Experimental Station (USDA NIFA), and a sponsored research agreement from VIRIS Detection Systems. Molecular graphics and analyses performed with UCSF ChimeraX, developed by the Resource for Biocomputing, Visualization, and Informatics at the University of California, San Francisco, with support from National Institutes of Health R01-GM129325 and the Office of Cyber Infrastructure and Computational Biology, National Institute of Allergy and Infectious Diseases. Funders had no role in designing, performing, interpreting, or submitting the work.

## Author Contributions

A.S.-F.: Conceptualization, Data Curation, Formal Analysis, Funding acquisition, Investigation, Methodology, Project administration, Resources, Supervision, Validation, Visualization, Writing – original draft. W.S.H., T.W., M.B.: Data curation, Investigation, Methodology, Visualization, Writing – review & editing. A.B.G., R.A.W.: Investigation, Methodology. L.T.: Visualization. C.C.G.: Software, Resources, Writing – review & editing. K.N. and E.E.: Investigation, Resources. G.C.L.: Methodology, Supervision, Visualization, Writing – review & editing. B.W.: Funding acquisition, Project administration, Resources, Supervision, Visualization, Writing – review & editing.

## Additional Information

**Supplementary Information** The online version contains supplementary material available at XXXXX.

**Supplementary Video 1**: Cas1-2/3 undergoes a large conformational change to unveil Cas2-leader binding sites.

**Supplementary Video 2**: Overview of how the I-F CRISPR leader and IHF guide Cas1-2/3-mediated integration of foreign DNA at the first CRISPR repeat.

**Supplementary Video 3**: The strained IHF-mediated DNA bends sway in relation to the Cas1-2/3 integrase.

**Correspondence and requests for materials** should be addressed to Blake Wiedenheft.

**Peer review information YYYY.**

**Reprints and permissions information** is available at http://www.nature.com/reprints.

## References

1. Koonin, E. V. & Krupovic, M. Evolution of adaptive immunity from transposable elements combined with innate immune systems. Nat. Rev. Genet. 16, 184–192 (2015).

2. McCLINTOCK, B. The origin and behavior of mutable loci in maize. Proc. Natl. Acad. Sci. U. S. A. 36, 344–355 (1950).

3. Nuñez, J. K., Bai, L., Harrington, L. B., Hinder, T. L. & Doudna, J. A. CRISPR Immunological Memory Requires a Host Factor for Specificity. Mol. Cell 62, 824–833 (2016).

4. Fagerlund, R. D. et al. Spacer capture and integration by a type I-F Cas1–Cas2-3 CRISPR adaptation complex. Proc. Natl. Acad. Sci. 114, 201618421 (2017).

5. Wright, A. V. et al. Structures of the CRISPR genome integration complex. Science 357, 1113–1118 (2017).

6. Hickman, A. B. & Dyda, F. Mechanisms of DNA transposition. Mob. DNA III 529–553 (2015) doi:10.1128/9781555819217.ch25.

7. Laxmikanthan, G. et al. Structure of a holliday junction complex reveals mechanisms governing a highly regulated DNA transaction. eLife 5, 1–23 (2016).

8. Lee, H. & Sashital, D. G. Creating memories: molecular mechanisms of CRISPR adaptation. Trends Biochem. Sci. 1–13 (2022) doi:10.1016/j.tibs.2022.02.004.

9. Wang, J. et al. Structural and Mechanistic Basis of PAM-Dependent Spacer Acquisition in CRISPR-Cas Systems. Cell 163, 840–853 (2015).

10. Nuñez, J. K., Harrington, L. B., Kranzusch, P. J., Engelman, A. N. & Doudna, J. A. Foreign DNA capture during CRISPR-Cas adaptive immunity. Nature 527, 535–538 (2015).

11. Xiao, Y., Ng, S., Nam, K. H. & Ke, A. How type II CRISPR–Cas establish immunity through Cas1–Cas2-mediated spacer integration. Nature 550, 137–141 (2017).

12. Jackson, S. A. et al. CRISPR-Cas: Adapting to change. Science 356, eaal5056 (2017).

13. Mojica, F. J. M., Díez-Villaseñor, C., García-Martínez, J. & Almendros, C. Short motif sequences determine the targets of the prokaryotic CRISPR defence system. Microbiology 155, 733–740 (2009).

14. Kim, S. et al. Selective loading and processing of prespacers for precise CRISPR adaptation. Nature 579, 141–145 (2020).

15. Hu, C. et al. Mechanism for Cas4-assisted directional spacer acquisition in CRISPR–Cas. Nature 598, 515– 520 (2021).

16. Ramachandran, A., Summerville, L., Learn, B. A., DeBell, L. & Bailey, S. Processing and integration of functionally oriented prespacers in the Escherichia coli CRISPR system depends on bacterial host exonucleases. J. Biol. Chem. 295, 3403–3414 (2020).

17. Liao, C. et al. Spacer prioritization in CRISPR–Cas9 immunity is enabled by the leader RNA. Nat. Microbiol. (2022) doi:10.1038/s41564-022-01074-3.

18. McGinn, J. & Marraffini, L. A. CRISPR-Cas Systems Optimize Their Immune Response by Specifying the Site of Spacer Integration. Mol. Cell 64, 616–623 (2016).

19. Wang, R., Li, M., Gong, L., Hu, S. & Xiang, H. DNA motifs determining the accuracy of repeat duplication during CRISPR adaptation in Haloarcula hispanica. Nucleic Acids Res. 44, 4266–4277 (2016).

20. Goren, M. G. et al. Repeat Size Determination by Two Molecular Rulers in the Type I-E CRISPR Array. Cell Rep. 16, 2811–2818 (2016).

21. Linheiro, R. S. & Bergman, C. M. Testing the palindromic target site model for DNA transposon insertion using the Drosophila melanogaster P-element. Nucleic Acids Res. 36, 6199–6208 (2008).

22. Santiago-Frangos, A., Buyukyoruk, M., Wiegand, T., Krishna, P. & Wiedenheft, B. Distribution and phasing of sequence motifs that facilitate CRISPR adaptation. Curr. Biol. 1–10 (2021) doi:10.1016/j.cub.2021.05.068.

23. Kieper, S. N., Almendros, C. & Brouns, S. J. J. Conserved motifs in the CRISPR leader sequence control spacer acquisition levels in Type I-D CRISPR-Cas systems. FEMS Microbiol. Lett. 366, 2016–2020 (2019).

24. Rollie, C., Graham, S., Rouillon, C. & White, M. F. Prespacer processing and specific integration in a Type I-A CRISPR system. Nucleic Acids Res. 46, 1007–1020 (2018).

25. Yosef, I., Goren, M. G. & Qimron, U. Proteins and DNA elements essential for the CRISPR adaptation process in Escherichia coli. Nucleic Acids Res. 40, 5569–5576 (2012).

26. Wei, Y., Chesne, M. T., Terns, R. M. & Terns, M. P. Sequences spanning the leader-repeat junction mediate CRISPR adaptation to phage in Streptococcus thermophilus. Nucleic Acids Res. 43, 1749–1758 (2015).

27. Wright, A. V. & Doudna, J. A. Protecting genome integrity during CRISPR immune adaptation. Nat. Struct. Mol. Biol. 23, 876–883 (2016).

28. Westra, E. R. et al. Parasite Exposure Drives Selective Evolution of Constitutive versus Inducible Defense. Curr. Biol. 25, 1043–1049 (2015).

29. Makarova, K. S. et al. Evolutionary classification of CRISPR–Cas systems: a burst of class 2 and derived variants. Nat. Rev. Microbiol. 18, 67–83 (2020).

30. Richter, C. et al. Priming in the Type I-F CRISPR-Cas system triggers strand-independent spacer acquisition, bi-directionally from the primed protospacer. Nucleic Acids Res. 42, 8516–8526 (2014).

31. Datsenko, K. A. et al. Molecular memory of prior infections activates the CRISPR/Cas adaptive bacterial immunity system. Nat. Commun. 3, 945 (2012).

32. Xiao, Y. et al. Structure basis for RNA-guided DNA degradation by Cascade and Cas3. 0839, 1–12 (2018).

33. Nicholson, T. J. et al. Bioinformatic evidence of widespread priming in type I and II CRISPR-Cas systems. RNA Biol. 16, 566–576 (2019).

34. Brown, M. W. et al. Assembly and translocation of a CRISPR-Cas primed acquisition complex. bioRxiv 41, 1–11 (2017).

35. Li, M., Wang, R., Zhao, D. & Xiang, H. Adaptation of the Haloarcula hispanica CRISPR-Cas system to a purified virus strictly requires a priming process. Nucleic Acids Res. 42, 2483–2492 (2014).

36. Semenova, E. et al. Highly efficient primed spacer acquisition from targets destroyed by the *Escherichia coli* type I-E CRISPR-Cas interfering complex. Proc. Natl. Acad. Sci. 113, 7626–7631 (2016).

37. Fineran, P. C. et al. Degenerate target sites mediate rapid primed CRISPR adaptation. Proc. Natl. Acad. Sci. U. S. A. 111, (2014).

38. Rice, P. A., Yang, S., Mizuuchi, K. & Nash, H. A. Crystal Structure of an IHF-DNA Complex: A Protein-Induced DNA U-Turn. Cell 87, 1295–1306 (1996).

39. Rohs, R. et al. Origins of specificity in protein-DNA recognition. Annu. Rev. Biochem. 79, 233–269 (2010).

40. Zayed, H. The DNA-bending protein HMGB1 is a cellular cofactor of Sleeping Beauty transposition. Nucleic Acids Res. 31, 2313–2322 (2003).

41. Little, A. J., Corbett, E., Ortega, F. & Schatz, D. G. Cooperative recruitment of HMGB1 during V(D)J recombination through interactions with RAG1 and DNA. Nucleic Acids Res. 41, 3289–3301 (2013).

42. Nash, H. A. & Robertson, C. A. Purification and properties of the Escherichia coli protein factor required for lambda integrative recombination. J. Biol. Chem. 256, 9246–9253 (1981).

43. Lavoie, B. D. & Chaconas, G. Site-specific HU binding in the Mu transpososome: conversion of a sequence-independent DNA-binding protein into a chemical nuclease. Genes Dev. 7, 2510–2519 (1993).

44. Chalmers, R., Guhathakurta, A., Benjamin, H. & Kleckner, N. IHF Modulation of Tn10 Transposition: Sensory Transduction of Supercoiling Status via a Proposed Protein/DNA Molecular Spring. Cell 93, 897– 908 (1998).

45. Haniford, D. B. Transpososome Dynamics and Regulation in Tn10 Transposition. Crit. Rev. Biochem. Mol. Biol. 41, 407–424 (2006).

46. Whitfield, C. R., Wardle, S. J. & Haniford, D. B. The global bacterial regulator H-NS promotes transpososome formation and transposition in the Tn5 system. Nucleic Acids Res. 37, 309–321 (2009).

47. Liu, D., Haniford, D. B. & Chalmers, R. M. H-NS mediates the dissociation of a refractory protein-DNA complex during Tn10/IS10 transposition. Nucleic Acids Res. 39, 6660–6668 (2011).

48. van Gent, D. C., Hiom, K., Paull, T. T. & Gellert, M. Stimulation of V(D)J cleavage by high mobility group proteins. EMBO J. 16, 2665–2670 (1997).

49. Rowland, S.-J., Stark, W. M. & Boocock, M. R. Sin recombinase from Staphylococcus aureus: synaptic complex architecture and transposon targeting: Sin recombinase. Mol. Microbiol. 44, 607–619 (2002).

50. Alonso, J. C., Weise, F. & Rojo, F. The Bacillus subtilis Histone-like Protein Hbsu Is Required for DNA Resolution and DNA Inversion Mediated by the β Recombinase of Plasmid pSM19035. J. Biol. Chem. 270, 2938–2945 (1995).

51. Petit, M.-A., Ehrlich, D. & Jannière, L. pAMβ1 resolvase has an atypical recombination site and requires a histone-like protein HU. Mol. Microbiol. 18, 271–282 (1995).

52. Rojo, F. & Alonso, J. C. The β recombinase of plasmid pSM19035 binds to two adjacent sites, making different contacts at each of them. Nucleic Acids Res. 23, 3181–3188 (1995).

53. Walker, M. W. G., Klompe, S. E., Zhang, D. J. & Sternberg, S. H. Transposon mutagenesis libraries reveal novel molecular requirements during CRISPR RNA-guided DNA integration. http://biorxiv.org/lookup/doi/10.1101/2023.01.19.524723 (2023) doi:10.1101/2023.01.19.524723.

54. Rollins, M. F. et al. Cas1 and the Csy complex are opposing regulators of Cas2/3 nuclease activity. Proc. Natl. Acad. Sci. 114, 201616395 (2017).

55. Wang, X. et al. Structural basis of Cas3 inhibition by the bacteriophage protein AcrF3. Nat. Struct. Mol. Biol. 23, 868–870 (2016).

56. Wiedenheft, B. et al. Structural Basis for DNase Activity of a Conserved Protein Implicated in CRISPR-Mediated Genome Defense. Structure 17, 904–912 (2009).

57. Kunin, V., Sorek, R. & Hugenholtz, P. Evolutionary conservation of sequence and secondary structures in CRISPR repeats. Genome Biol. 8, R61 (2007).

58. Nethery, M. A. et al. CRISPRclassify: Repeat-Based Classification of CRISPR Loci. CRISPR J. 4, 558–574 (2021).

59. Dhingra, Y., Suresh, S. K., Juneja, P. & Sashital, D. G. PAM binding ensures orientational integration during Cas4-Cas1-Cas2-mediated CRISPR adaptation. Mol. Cell 82, 4353–4367.e6 (2022).

60. Ali Azam, T., Iwata, A., Nishimura, A., Ueda, S. & Ishihama, A. Growth Phase-Dependent Variation in Protein Composition of the *Escherichia coli* Nucleoid. J. Bacteriol. 181, 6361–6370 (1999).

61. Montaño, S. P., Pigli, Y. Z. & Rice, P. A. The Mu transpososome structure sheds light on DDE recombinase evolution. Nature 491, 413–417 (2012).

62. Maertens, G. N., Hare, S. & Cherepanov, P. The mechanism of retroviral integration from X-ray structures of its key intermediates. Nature 468, 326–329 (2010).

63. Rollie, C., Schneider, S., Brinkmann, A. S., Bolt, E. L. & White, M. F. Intrinsic sequence specificity of the Cas1 integrase directs new spacer acquisition. eLife 4, 1–19 (2015).

64. Makarova, K. S., Wolf, Y. I. & Koonin, E. V. Classification and Nomenclature of CRISPR-Cas Systems: Where from Here? CRISPR J. 1, 325–336 (2018).

65. Deveau, H. et al. Phage response to CRISPR-encoded resistance in Streptococcus thermophilus. J. Bacteriol. 190, 1390–1400 (2008).

66. Künne, T. et al. Cas3-Derived Target DNA Degradation Fragments Fuel Primed CRISPR Adaptation. Mol. Cell 63, 852–864 (2016).

67. Musharova, O., et al. Prespacers formed during primed adaptation associate with the Cas1–Cas2 adaptation complex and the Cas3 interference nuclease–helicase. Proc. Natl. Acad. Sci. 118, e2021291118 (2021).

68. Wiegand, T. et al. Reproducible Antigen Recognition by the Type I-F CRISPR-Cas System. CRISPR J. 3, 378–387 (2020).

69. Vorontsova, D. et al. Foreign DNA acquisition by the I-F CRISPR–Cas system requires all components of the interference machinery. Nucleic Acids Res. 43, 10848–10860 (2015).

70. Koonin, E. V. & Makarova, K. S. Evolutionary plasticity and functional versatility of CRISPR systems. PLOS Biol. 20, e3001481 (2022).

71. Cavazzana-Calvo, M. et al. Gene Therapy of Human Severe Combined Immunodeficiency (SCID)-X1 Disease. Science 288, 669–672 (2000).

72. Strecker, J. et al. RNA-guided DNA insertion with CRISPR-associated transposases. Science 365, 48–53 (2019).

73. Klompe, S. E., Vo, P. L. H., Halpin-Healy, T. S. & Sternberg, S. H. Transposon-encoded CRISPR–Cas systems direct RNA-guided DNA integration. Nature 571, 219–225 (2019).

74. Shipman, S. L., Nivala, J., Macklis, J. D. & Church, G. M. Molecular recordings by directed CRISPR spacer acquisition. Science 353, aaf1175 (2016).

75. Schmidt, F., Cherepkova, M. Y. & Platt, R. J. Transcriptional recording by CRISPR spacer acquisition from RNA. Nature 562, 380–385 (2018).

76. Herzik, M. A., Wu, M. & Lander, G. C. High-resolution structure determination of sub-100 kDa complexes using conventional cryo-EM. Nat. Commun. 10, 1–9 (2019).

77. Mastronarde, D. N. Automated electron microscope tomography using robust prediction of specimen movements. J. Struct. Biol. 152, 36–51 (2005).

78. Punjani, A., Rubinstein, J. L., Fleet, D. J. & Brubaker, M. A. CryoSPARC: Algorithms for rapid unsupervised cryo-EM structure determination. Nat. Methods 14, 290–296 (2017).

79. Suloway, C. et al. Automated molecular microscopy: The new Leginon system. J. Struct. Biol. 151, 41–60 (2005).

80. Punjani, A., Zhang, H. & Fleet, D. J. Non-uniform refinement: adaptive regularization improves single-particle cryo-EM reconstruction. Nat. Methods 17, 1214–1221 (2020).

81. Scheres, S. H. W. & Chen, S. Prevention of overfitting in cryo-EM structure determination. Nat. Methods 9, 853–854 (2012).

82. Tan, Y. Z. et al. Addressing preferred specimen orientation in single-particle cryo-EM through tilting. Nat. Methods 14, 793–796 (2017).

83. Liebschner, D. et al. Macromolecular structure determination using X-rays, neutrons and electrons: recent developments in *Phenix*. Acta Crystallogr. Sect. Struct. Biol. 75, 861–877 (2019).

84. Mirdita, M. et al. ColabFold: making protein folding accessible to all. Nat. Methods 19, 679–682 (2022).

85. Emsley, P., Lohkamp, B., Scott, W. G. & Cowtan, K. Features and development of Coot. Acta Crystallogr. Sect. D 66, 486–501 (2010).

86. Nicholls, R. A. Conformation-independent comparison of protein structures. (2011).

87. Williams, C. J. et al. MolProbity: More and better reference data for improved all-atom structure validation: PROTEIN SCIENCE.ORG. Protein Sci. 27, 293–315 (2018).

88. Goddard, T. D. et al. UCSF ChimeraX: Meeting modern challenges in visualization and analysis. Protein Sci. 27, 14–25 (2018).

89. Pettersen, E. F. et al. UCSF ChimeraX: Structure visualization for researchers, educators, and developers. Protein Sci. 30, 70–82 (2021).

90. Sagendorf, J. M., Markarian, N., Berman, H. M. & Rohs, R. DNAproDB: an expanded database and web-based tool for structural analysis of DNA–protein complexes. Nucleic Acids Res. gkz889 (2019) doi:10.1093/nar/gkz889.

91. Biswas, A., Staals, R. H. J., Morales, S. E., Fineran, P. C. & Brown, C. M. CRISPRDetect: A flexible algorithm to define CRISPR arrays. BMC Genomics 17, 356 (2016).

92. Hyatt, D. et al. Prodigal: prokaryotic gene recognition and translation initiation site identification. BMC Bioinformatics 11, 119 (2010).

93. Abby, S. S., Néron, B., Ménager, H., Touchon, M. & Rocha, E. P. C. MacSyFinder: A program to mine genomes for molecular systems with an application to CRISPR-Cas systems. PLoS ONE (2014) doi:10.1371/journal.pone.0110726.

94. Couvin, D. et al. CRISPRCasFinder, an update of CRISRFinder, includes a portable version, enhanced performance and integrates search for Cas proteins. Nucleic Acids Res. 46, W246–W251 (2018).

95. Li, W. & Godzik, A. Cd-hit: a fast program for clustering and comparing large sets of protein or nucleotide sequences. Bioinformatics 22, 1658–1659 (2006).

96. Grant, C. E., Bailey, T. L. & Noble, W. S. FIMO: scanning for occurrences of a given motif. Bioinformatics 27, 1017–1018 (2011).

97. Gouveia-Oliveira, R., Sackett, P. W. & Pedersen, A. G. MaxAlign: maximizing usable data in an alignment. BMC Bioinformatics 8, 312 (2007).

98. Finn, R. D., Clements, J. & Eddy, S. R. HMMER web server: interactive sequence similarity searching. Nucleic Acids Res. 39, W29–W37 (2011).

99. Suzek, B. E. et al. UniRef clusters: a comprehensive and scalable alternative for improving sequence similarity searches. Bioinformatics 31, 926–932 (2015).

100. Katoh, K. & Standley, D. M. MAFFT Multiple Sequence Alignment Software Version 7: Improvements in Performance and Usability. Mol. Biol. Evol. 30, 772–780 (2013).

101. Schneider, T. D. & Stephens, R. M. Sequence logos: A new way to display consensus sequences. Nucleic Acids Res. 18, 6097–6100 (1990).

102. Crooks, G. E., Hon, G., Chandonia, J.-M. & Brenner, S. E. WebLogo: a sequence logo generator. Genome Res. 14, 1188–90 (2004).

