## Extended Data for "Protein-mediated genome folding allosterically enhances site-specific integration of foreign DNA into CRISPRs"

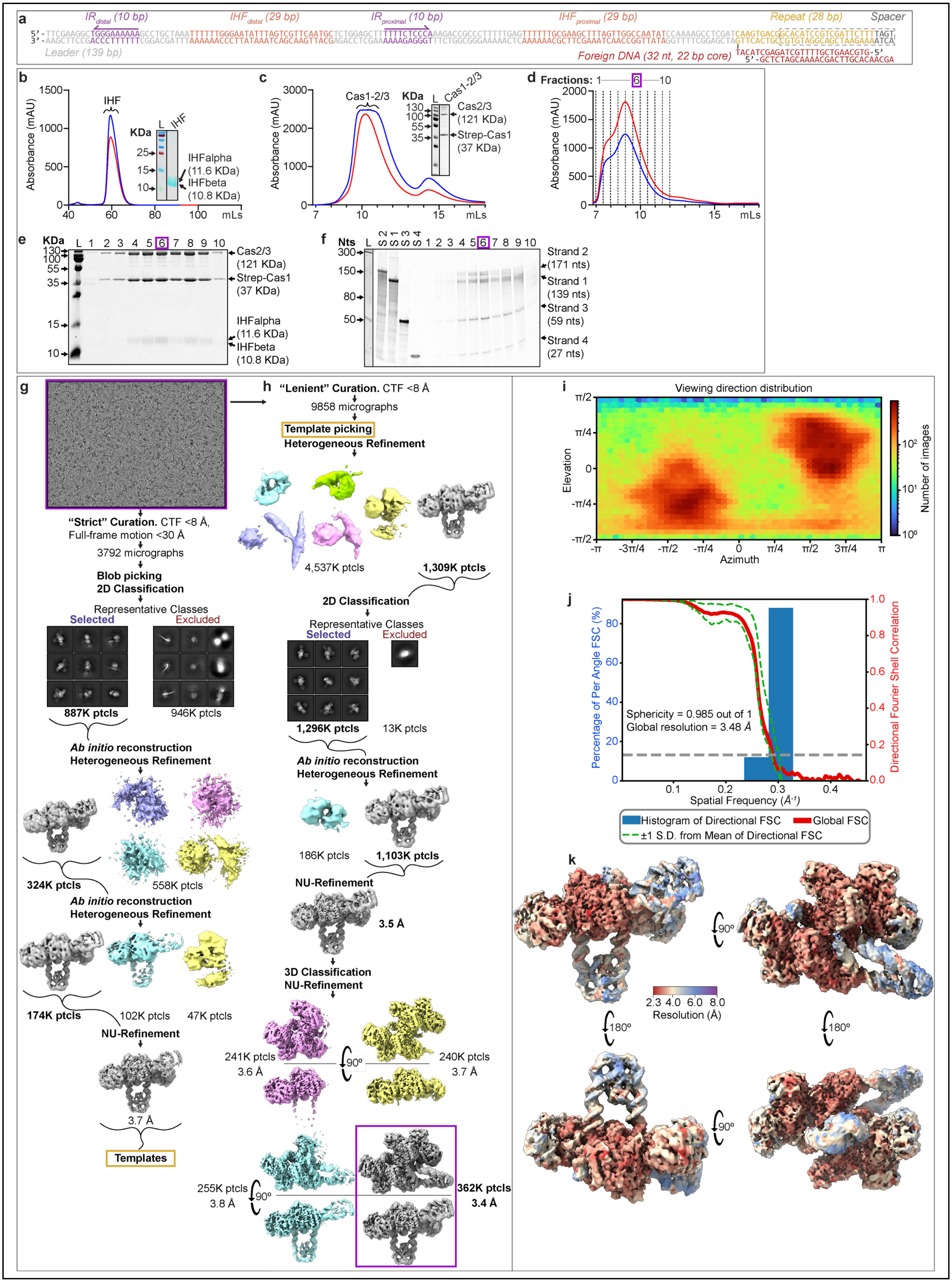


**Extended Data Fig. 1 Cryo-EM sample preparation, imaging and processing for type I-F integration complex. a,**

Sequence-level schematic of DNA used to assemble the integration complex. The length of each motif is listed. The latter two thirds of the CRISPR repeat and second spacer (grey dashed box) could not be resolved in the cryo-EM reconstruction. See also **Supplementary Table S1**. **b**, Size-exclusion chromatography (SEC) profile (Superdex 75 16/600, Cytiva) of IHF heterodimer purified as described in methods section, and SDS-PAGE gel (inset). **c**, SEC profile (Superdex 200 10/300, Cytiva) of Cas1-2/3 heterohexamer purified as described in methods section, and SDS-PAGE gel (inset). **d**, I-F integration complex was assembled from purified DNAs, IHF and Cas1-2/3 as described in the methods section, and the assembled complex was further purified by size-exclusion chromatography (SEC) (Superdex 200 10/300, Cytiva). Individual fractions were collected along the elution profile, and were concentrated and stored separately for further analysis and imaging. **e**, Individual SEC fractions were analyzed by SDS-PAGE to determine which fractions contained all the proteins necessary for a complete complex. **f**, Individual SEC fractions were phenol-chloroform extracted, and the aqueous layer was analyzed by Urea-PAGE to determine which fractions contained all four DNA strands necessary for a complete complex. The fraction chosen for cryo-EM analysis is indicated with a dotted purple box. **g**, Image processing pipeline for a small subset of 10,740 total micrographs for the type I-F integration complex, to generate an initial model for template picking. **h**, Final image processing pipeline for the type I-F integration complex. **i**, Viewing direction distribution plot depicting particle orientations present in final reconstruction. More populated views are shown in red, and less populated views are shown in blue. **j**, 3D Fourier Shell Correlation (3DFSC) of the final I-F integration complex reconstruction. The global resolution at 0.143 is indicated by a dashed line, 3.48Å. **k**, Local resolution estimation of the cryo-EM reconstruction calculated by cryoSPARC[1].

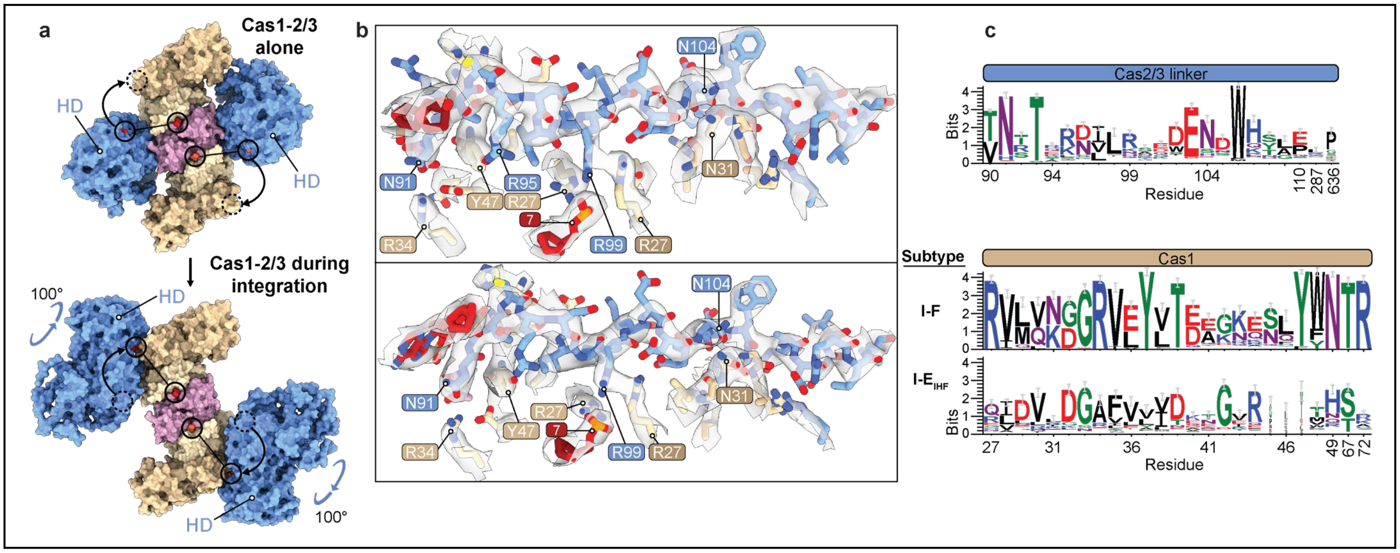


**Extended Data Fig. 2 Cas1-2/3 undergoes a large structural rearrangement during integration. a**, The Cas3 domains of the Cas1-2/3 complex have undergone a ~100° rotation in the structure of the integration complex as compared to Cas1-2/3 alone [2]. The positions of the first (90) and last residues (110) of the Cas2/3 linker are shown (Cas2/3a, red; Cas2/3b, tomato). The linker residues were not resolved for the previously determined pseudo-atomic model of Cas1-2/3 alone, and so are not shown for either complex for clarity[2]. The rotation of the Cas3 domains outwards unveils new DNA binding sites on two opposing faces of the Cas2 dimer. **b**, Zoom-in on the atomic fit of the Cas2/3 linker to the cryo-EM map. The Cas2/3 linker is disordered in the absence of Cas1 (PDB:5B7I) [3], but has become ordered in the I-F integration complex structure due to packing by the foreign DNA against the Cas1 beta hairpins. **c**, A sequence logo depicting the conservation of Cas2/3 linker residues (top). The Cas1 residues that contact the Cas2/3 linker are conserved in Cas1 proteins associated with type I-F CRISPR loci (middle), but are not conserved in the closely related Cas1 proteins associated with type I-E CRISPR loci that have similar IHF motif-containing leaders [4]. Residues are numbered according to *P. aeruginosa* PA14 Cas1 and Cas2/3 proteins.

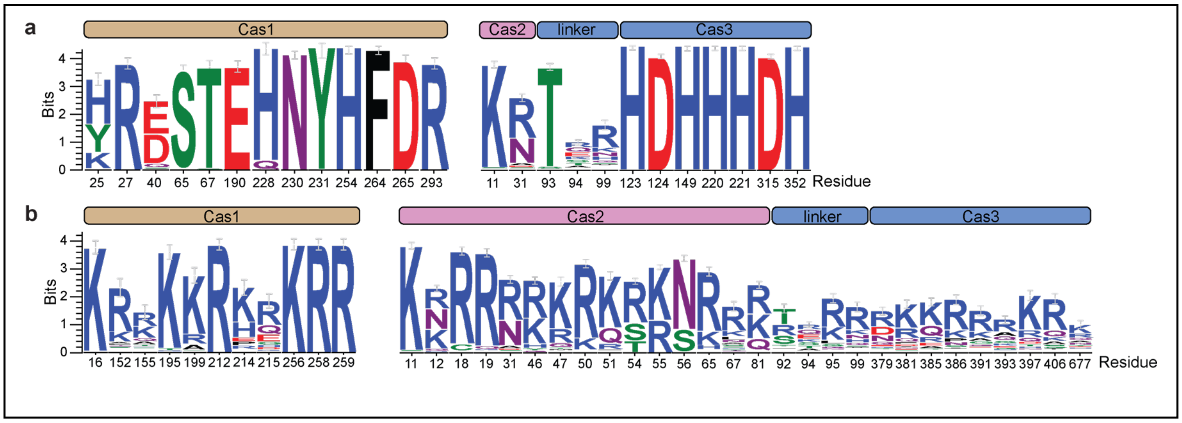


**Extended Data Fig. 3 Conservation analysis of Cas1-2/3 residues involved in DNA binding and integration. a**, Conservation of Cas1 and Cas2/3 residues involved in binding the foreign DNA, or catalyzing the strand transfer reaction, or catalyzing the degradation of nucleic acids. See Figure 2. **b**, Conservation of basic and polar Cas1 and Cas2/3 residues involved in accommodating the DNA duplexes bound by the Cas1-2/3 complex during integration. See Figure 3. Residues are numbered according to *P. aeruginosa* Cas1 and Cas2/3 proteins.

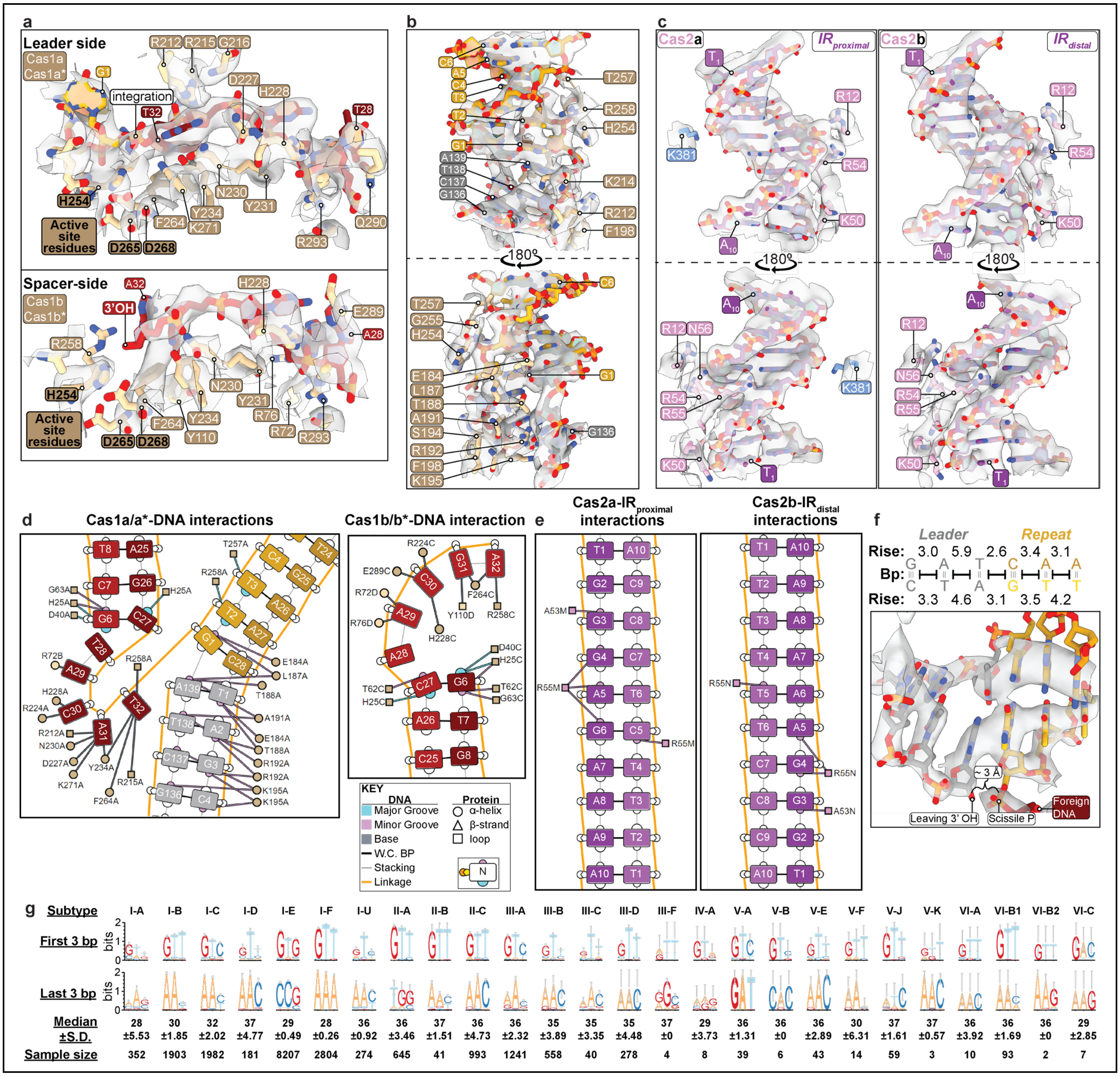


**Extended Data Fig. 4 Cas1-2/3 predominantly recognizes IR motifs, CRISPR repeat and foreign DNA through non-sequence specific interactions. a**, Splayed 3' ends of the foreign DNA are directed into the Cas1 transesterification active site. The product of the first strand-transfer reaction is shown in the Cas1a* active site (top), and the 3' OH of the other end of the foreign DNA is positioned in the Cas1b* active site (bottom). The cryo-EM map is shown in transparent grey. **b**, Zoom-in on the Cas1-2/3 contacts to the CRISPR repeat (ChimeraX *contacts* command with default parameters). Most protein contacts occur to the DNA backbone and minor groove. Cas1 residue E184 appears to probe nucleotide G1 of the repeat. **c**, Zoom-in on the Cas1-2/3 contacts to the IR leader motifs. Most protein contacts occur to the DNA backbone and minor groove. **d**, DNAproDB analysis of Cas1-2/3 interactions with the 3' ends of foreign DNA and the CRISPR leader-repeat junction. For clarity, only protein interactions to the nucleobases are shown[5]. **e**, DNAproDB analysis of Cas1-2/3 interactions with the IR leader motifs [5]. For clarity, only protein interactions to the nucleobases are shown. **f**, Zoom-in on the atomic fit of the base-pairs around the leader-repeat junction (-3 and +3 bps, coordinated by Cas1a*), to the cryo-EM map. Tension in the DNA loop at the leader-repeat junction has been released in the post-integration structure by a physical separation of base-pairs, as measured by an increase in base step rise. This tension may further pull the leaving 3'OH out of the Cas1 transesterification active site, to inhibit disintegration of the foreign DNA from the repeat. The approximate local base step rise was calculated using the http://web.x3dna.org/ webserver. **g**, A bioinformatic analysis of the first repeat from 24,940 CRISPR loci reveals that a 5' GT dinucleotide is strongly conserved across most CRISPR subtypes. Similarly, a 5' GT is present at the spacer-end of the repeat (seen as AC-3' on the sense strand) within certain CRISPR subtypes (I-D, II-C, III-C, III-D, V-B, V-E, V-K, VI-A), but it is not broadly conserved.

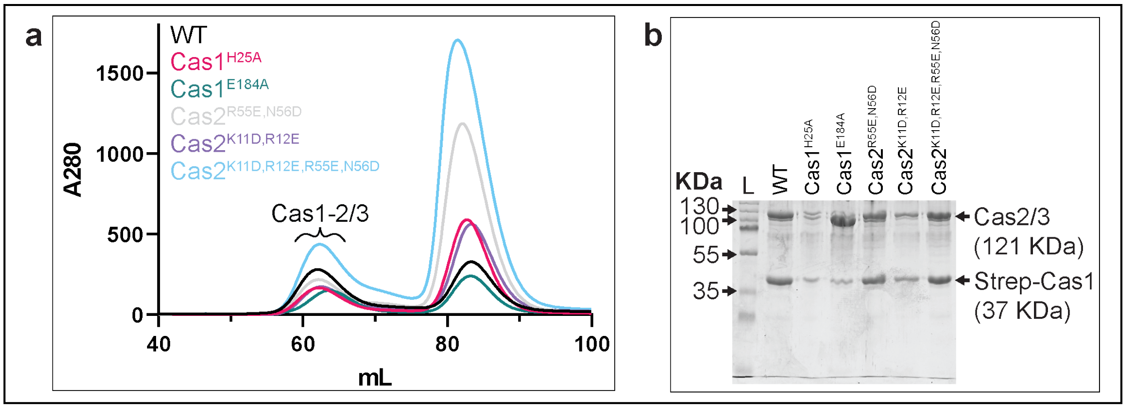


**Extended Data Fig. 5 Purification of structure-guided mutants of Cas1-2/3. a**, SEC profile of a new preparation of wildtype Cas1-2/3 and all variants purified in the same manner on a Superdex 200 16/600 (Cytiva). An excess of free Strep-tagged Cas1 elutes at approximately 82 mL. **b**, SDS-PAGE gel of the Cas1-2/3 hetero-hexamer peak for all purified Cas1-2/3 variants.

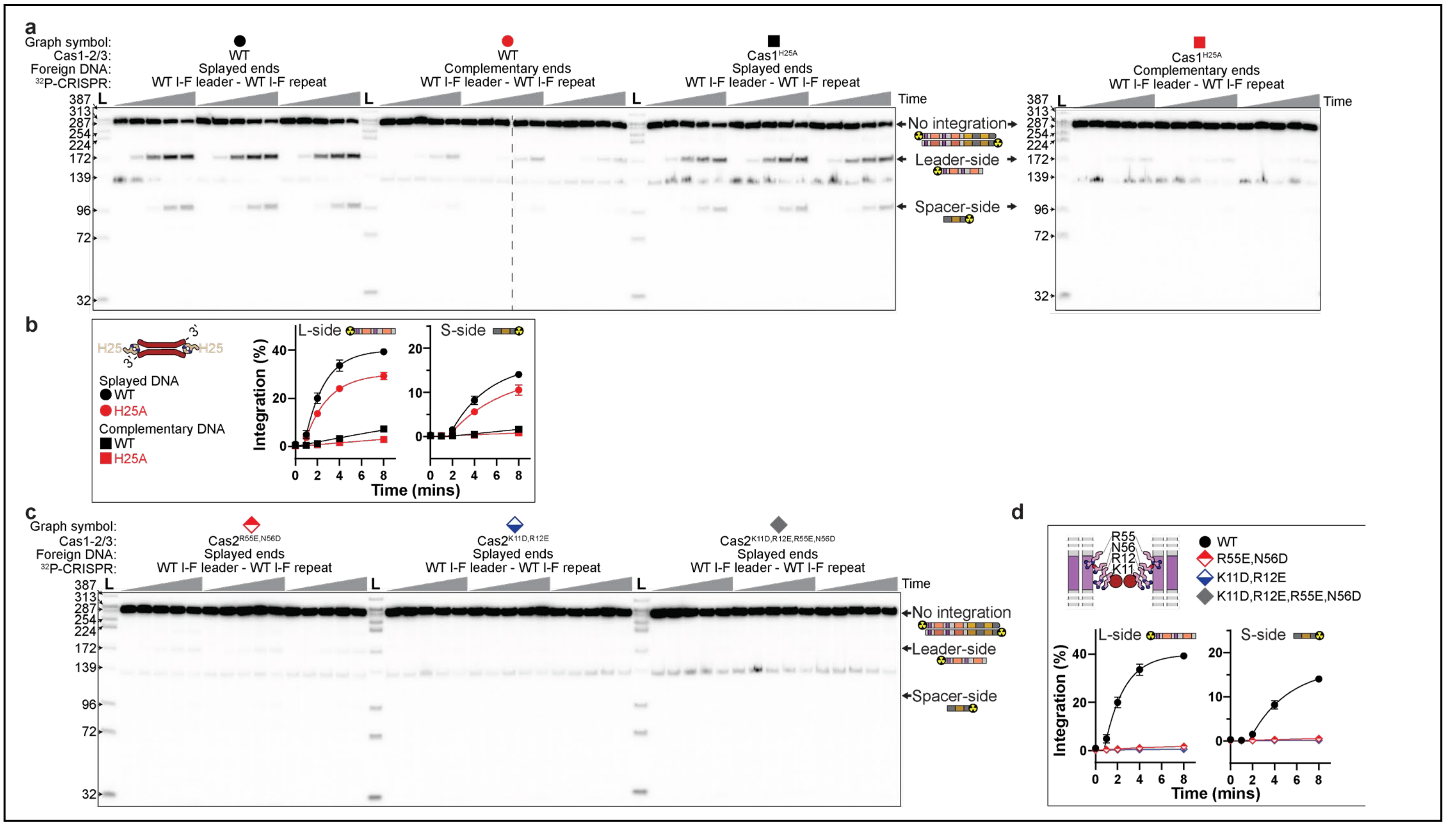


**Extended Data Fig. 6 Validation of Cas1-2/3 interactions with the foreign DNA and IR motifs. a**, Time-course integration reactions to test the role of Cas1$^{H25}$ in splaying the foreign DNA ends. Integration reactions were performed with trimmed foreign DNA (lacking a PAM) in triplicate, resolved on denaturing polyacrylamide gels. Timepoints were taken at 0, 1, 2, 4 and 8 minutes. Reactions were stopped by the addition of phenol. A $^{32}$P-labelled DNA that is shorter (140-160 bp) than the full length CRISPR is present in some DNA preparations (also see Extended Data Fig. 6c). Full-length CRISPR DNA, leader- and spacer-side integration products, do not overlap with this band. Further, Cas1-2/3, foreign DNA and IHF are in excess over the 32P-labelled DNA. The 140-160 bp band does not interfere with the quantification or generation of integration products. **b**, Quantification of time-course experiments to determine the role of Cas1 residue H25 in integration. The Cas1$^{H25A}$ mutant integrates splayed and fully complementary foreign DNA fragments less efficiently that WT Cas1-2/3, suggesting that H25 steers the non-nucleophilic DNA strand away from the Cas1 active site. These results mirror the previously published effect of type I-E Cas1-2 tyrosine wedge mutation [6]. **c**, Time-course integration reactions to test the role of Cas2 residues in recognition of the IR motifs in the leader, performed as in panel a. **d**, Quantification of time-course experiments to determine the role of Cas2 residues K11, R12, R55 and N56 in integration. The Cas2$^{R55E,N56D}$/3 mutant retains a small amount of integration activity. The Cas2$^{K11D,R12E}$/3 and Cas2$^{K11D,R12E,R55E,N56D}$/3 mutants do not integrate DNA into the I-F CRISPR.

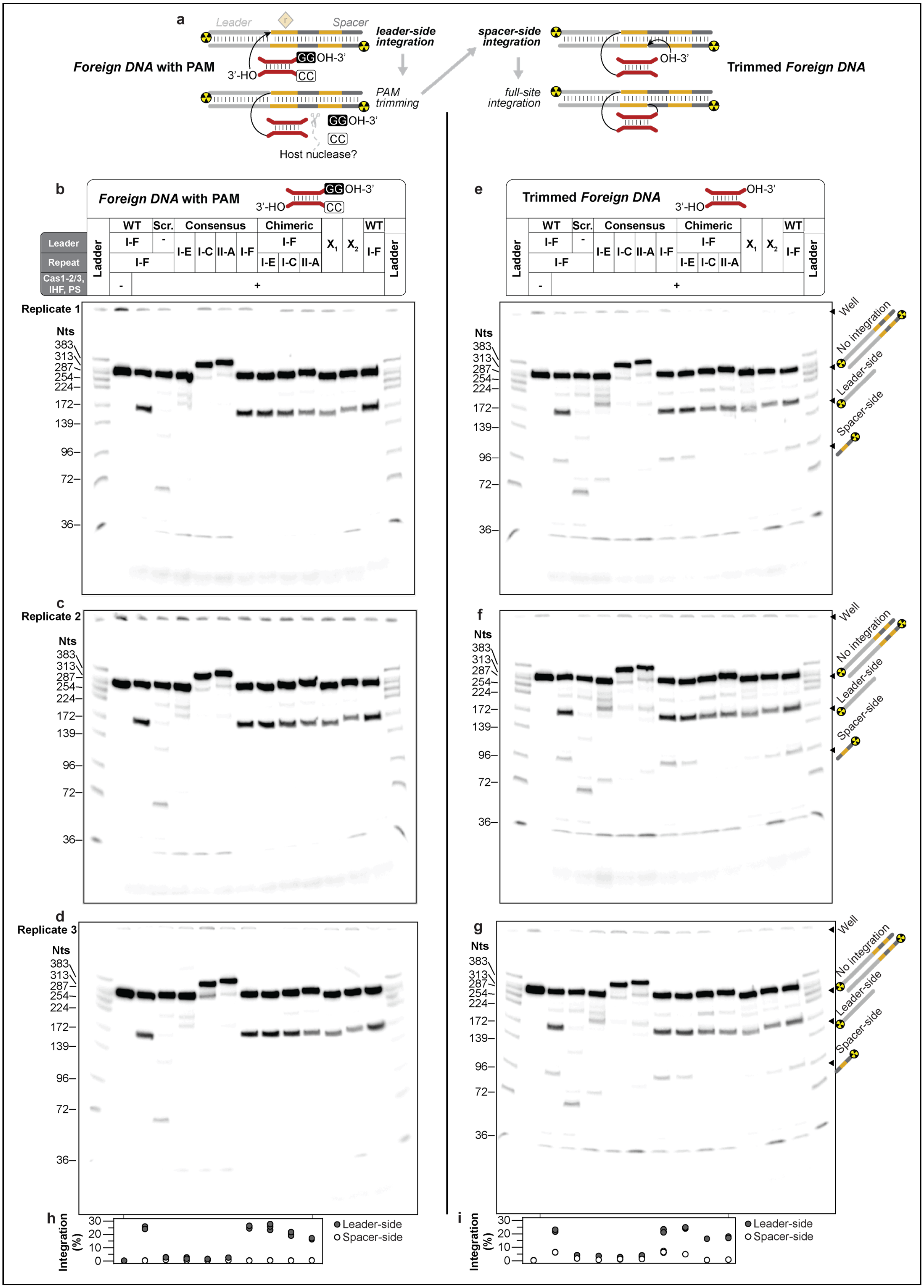
a
Leader
Spacer
Foreign DNA with PAM
leader-side integration
PAM trimming
Host nuclease?
spacer-side integration
full-site integration
Trimmed Foreign DNA
b
Foreign DNA with PAM
e
Trimmed Foreign DNA
Leader
Repeat
Cas1-2/3, IHF, PS
Ladder
WT
Scr.
Consensus
Chimeric
I-F
I-E
I-C
II-A
X1
X2
Replicate 1
Replicate 2
Replicate 3
Nts
c
d
f
g
Well
No integration
Leader-side
Spacer-side
h
i
Integration (%)
Leader-side
Spacer-side

**Extended Data Fig. 7 PAM blocks Cas-mediated integration of foreign DNA into CRISPR repeat. a**, Schematic summarizing the step-wise strand-transfer reactions catalyzed by a Cas integrase. In the absence of putative host nucleases that trim PAMs, the foreign DNA fragments stall the reaction at leader-side integration. However, the foreign DNA fragments that contain a PAM proceed through leader- and spacer-side integration. **b-d**, Endpoint integration reactions performed with a PAM-containing foreign DNA in triplicate, resolved on denaturing polyacrylamide gels. The X1 and X2 lanes signify lanes that were not further analyzed for this manuscript. **e-g**, Endpoint integration reactions performed with a trimmed foreign DNA in triplicate, resolved on denaturing polyacrylamide gels. The X1 and X2 lanes signify integration substrates that were not analyzed for this manuscript. **h, i**, Quantification of leader- (grey circles) or spacer-side (white circles) integration events from all three replicate gels. Individual dots for each triplicate reaction are shown, and some dots overlap.

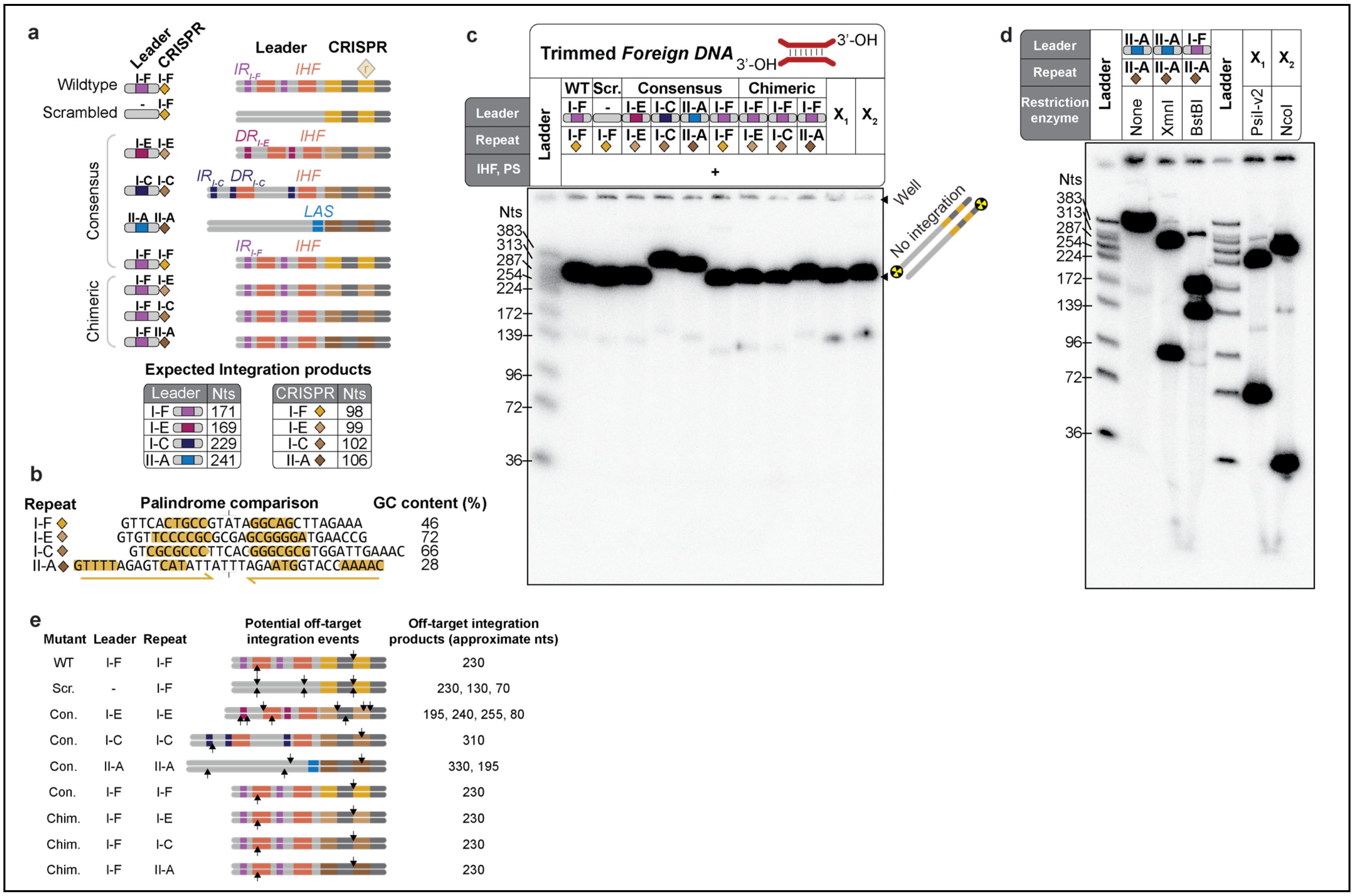


**Extended Data Fig. 8 Control reactions for integration assay and generation of $^{32}$P-labelled ladder. a**, Scheme of nine CRISPR fragments used for *in vitro* integration assays. Each CRISPR locus contains two repeats and two spacers. Leader motifs are color-coded and annotated (IR, inverted repeat; DR, direct repeat; IHF, IHF binding site; LAS, Leader anchoring site). To simplify the pictograms shown here and in subsequent panels, a single-colored rectangle was used to represent a given collection of leader motifs, and a single diamond was used to represent a CRISPR locus composed of two repeats and two spacers. **b**, The four CRISPR repeats tested in the integration assays have diverse palindromes (yellow), and a wide range of GC-content. **c**, Control reactions in which all components necessary for integration, except Cas1-2/3, were incubated. The overexposed gels show that the majority of the $^{32}$P signal for a given integration substrate DNA corresponds to the full-length strands. **d**, A custom $^{32}$P-labelled DNA ladder was made by mixing the degradation products generated by individual restriction enzyme digests of different $^{32}$P-labelled integration substrate DNAs. X1 and X2 signify integration substrates that were not analyzed for this manuscript. **e**, Schematics of the nine CRISPRs tested in Extended Data Fig. 5. The arrows identify locations of off-target integration reactions. Most off-target integration reactions occur by spurious integration at DNA motifs (e.g., second CRISPR repeat, IHF$^{distal}$ site, or upstream motifs) found near the ends of the CRISPR DNA target. Previous deep-sequencing of similar integration reactions has shown that the second repeat is a common off-target integration site, and that IHF blocks integration at the IHF binding site [4].

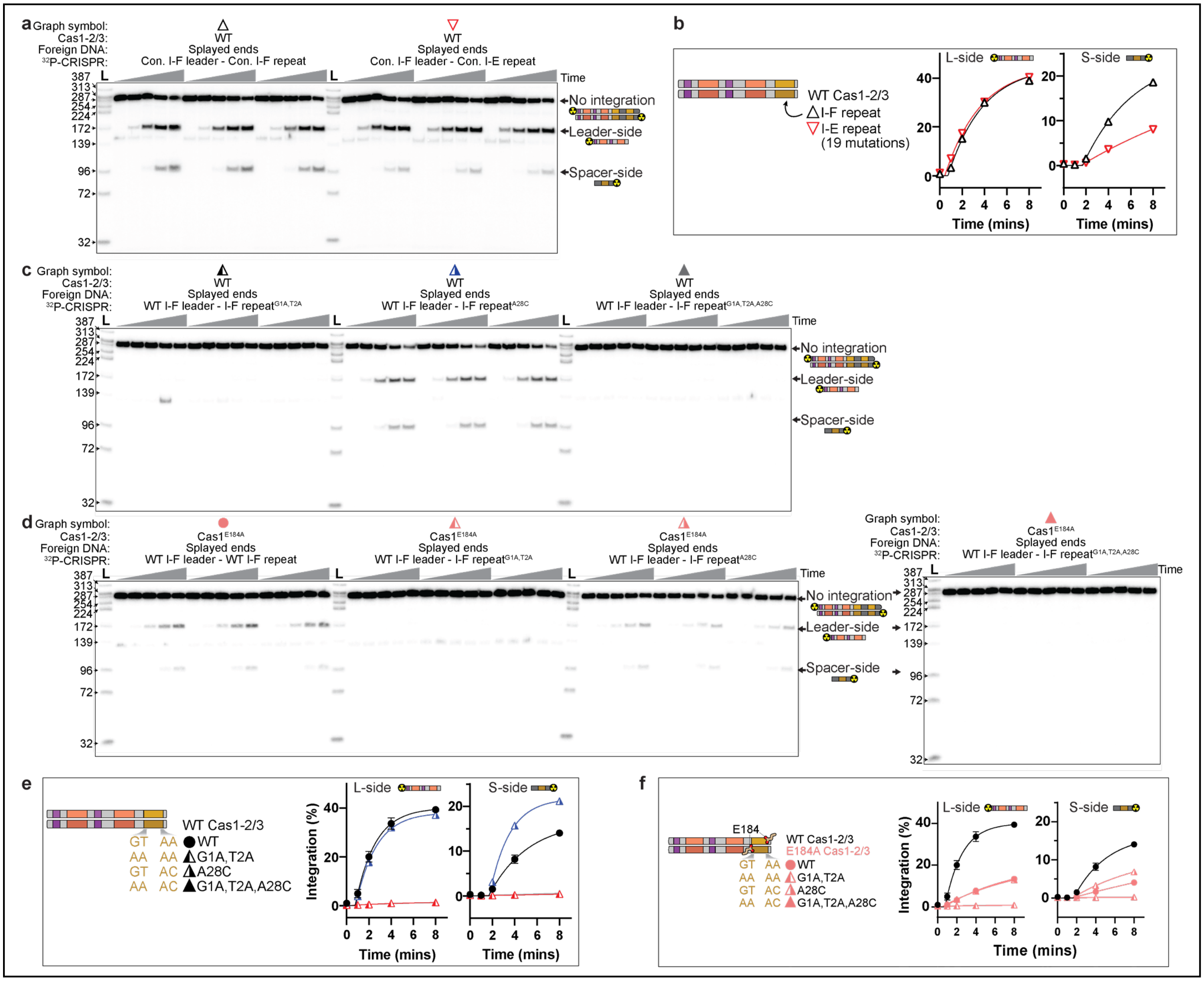


**Extended Data Fig. 9 Validation of Cas1-2/3 interactions with the repeat. a**, Time-course integration reactions to compare rate of integration into I-E and I-F repeats downstream of a I-F leader. **b**, Quantification of time-course experiments to determine the impact of 19 mutations associated with swapping the I-F repeat for the I-E repeat, on integration. Leader-side integration is indistinguishable. But spacer-side integration is slower into the I-E repeat. **c**, Time-course integration reactions to measure the impact of I-F repeat mutations on integration rate. **d**, Time-course integration reactions to measure the impact of I-F repeat mutations on integration rate, in the context of a Cas1$^{E184A}$ mutation. The Cas1$^{E184A}$ mutation is expected to disrupt 5' G recognition, but also impacts stability of the Cas1-2/3 complex (**Extended Data Fig. 5**). **e**, Quantification of time-course experiments to determine the impact of I-F repeat mutations on integration rate, in the context of WT Cas1-2/3. **f**, Quantification of time-course experiments to determine the impact of I-F repeat mutations on integration rate, in the context of Cas1$^{E184A}$-2/3.

### References


1. Punjani, A., Rubinstein, J. L., Fleet, D. J. & Brubaker, M. A. CryoSPARC: Algorithms for rapid unsupervised cryo-EM structure determination. *Nature Methods* **14**, 290–296 (2017).
2. Rollins, M. F. *et al.* Cas1 and the Csy complex are opposing regulators of Cas2/3 nuclease activity. *Proceedings of the National Academy of Sciences* **114**, 201616395 (2017).
3. Wang, X. *et al.* Structural basis of Cas3 inhibition by the bacteriophage protein AcrF3. *Nature Structural and Molecular Biology* **23**, 868–870 (2016).
4. Santiago-Frangos, A., Buyukyoruk, M., Wiegand, T., Krishna, P. & Wiedenheft, B. Distribution and phasing of sequence motifs that facilitate CRISPR adaptation. *Current Biology* 1–10 (2021) doi:10.1016/j.cub.2021.05.068.
5. Sagendorf, J. M., Markarian, N., Berman, H. M. & Rohs, R. DNAproDB: an expanded database and web-based tool for structural analysis of DNA–protein complexes. *Nucleic Acids Research* gkz889 (2019) doi:10.1093/nar/gkz889.
6. Nuñez, J. K., Harrington, L. B., Kranzusch, P. J., Engelman, A. N. & Doudna, J. A. Foreign DNA capture during CRISPR-Cas adaptive immunity. *Nature* **527**, 535–538 (2015).
