## Supplementary Table S1 for "Protein-mediated genome folding allosterically enhances site-specific integration of foreign DNA into CRISPRs"

**Supplementary Table 1. Oligonucleotides used in this study.**

| Name | Description | Sequence (5' to 3') |
|---|---|---|
| Strand_1 | I-F leader sense strand, for I-F complex assembly - ssDNA | AAGCTTCCGACCCTTTTTTCGGACGATTTAAAAAACCCTTATAAATC AGCAAGTTACGAGACCTCGAAAAAAGAGGGTTTCTGGCGGGAAAA ACTCAAAAAACGCTTCGAAATCAACCGGTTATAGGTTTTCGGAGCT A |
| Strand_2 | I-F leader and repeat anti-sense strand, for I-F complex assembly - ssDNA | TGATTTTCTTAGCTGCCTACACGGCAGTGAACTAGCTCCGAAAACC TATAACCGGTTGATTTCGAAGCGTTTTTTGAGTTTTTCCCGCCAGA AACCCTCTTTTTTCGAGGTCTCGTAACTTGCTGATTTATAAGGGTTT TTTAAATCGTCCGAAAAAAGGGTCGGAAGCTT |
| Strand_3 | I-F repeat and prespacer sense strand, for I-F complex assembly - ssDNA | GTGCAAGTCGTTTTGCTAGAGCTACATGTTCACTGCCGTGTAGGC AGCTAAGAAAATCA |
| Strand_4 | Foreign DNA anti-sense strand, for I-F complex assembly - ssDNA | GCTCTAGCAAAACGACTTGCACAACGA |
| Splayed, Trimmed foreign DNA | DNA with 22 bp core and 5 nt overhangs on all 5' and 3' ends -dsDNA, for integration assays | TACATGCTCTAGCAAAACGACTTGCACAACGA *and* ATTAAGTGCAAGTCGTTTTGCTAGAGCTACAT |
| Fully complementary, Trimmed foreign DNA | 32 bp DNA, for integration assays | TTAAT GCTCTAGCAAAACGACTTGCAC TTAAT *and* ATTAA GTGCAAGTCGTTTTGCTAGAGC ATTAA |
| Splayed, PAM-containing foreign DNA | DNA with 22 bp core, 5 nt overhangs on one set of ends, and 7 nts overhangs on the other end (which contains the PAM) – dsDNA, for integration assays | TACATGCTCTAGCAAAACGACTTGCACAACGAGG *and* CCATTAAGTGCAAGTCGTTTTGCTAGAGCTACAT |
| Wildtype I-F Leader and wildtype I-F Locus DNA | EcoRI and BamHI digest of Addgene plasmid 192207, $^{32}$P-labelled for integration assays | AATTCGAGCTCGGTACCTCGCGAATGCATCTAGATCCATGGACCC TTTTTTCGGACGATTTCTTACGCCCTTATAAATCAGCAAGTTACGA GACCTCGAAAAAAGAGGGTTTCTGGCGGGAAAAACTCGGTATTTC TTTTTCCTTCAAATGGTTATAGGTTTTCGGAGCTAGTTCACTGCCG TGTAGGCAGCTAAGAAAATCAGCCGGACGTTGTAGTAGTCGAGCG CGGTGTTCACTGCCGTGTAGGCAGCTAAGAAAGCCGGGTGGTGA CTTCTGGACGGCGTTCGTCAAGGATC |
| Scrambled I-F Leader and wildtype I-F Locus DNA | EcoRI and BamHI digest of Addgene plasmid 192208, $^{32}$P-labelled for integration assays | AATTCGAGCTCGGTACCTCGCGAATGCATCTAGATCCATGGTCTTA TCTCTCGGACGATTTTATTCTTGTACCACGTCACACGGTCAAAAAG ACCTCGAAGAGAGTGAAATTCTGGCGGGAAAAACTCTGTATTAACT CCGTTATGTATTTTTGACTGGTTTTCGGAGCTAGTTCACTGCCGTG TAGGCAGCTAAGAAAATCAGCCGGACGTTGTAGTAGTCGAGCGCG GTGTTCACTGCCGTGTAGGCAGCTAAGAAAGCCGGGTGGTGACTT CTGGACGGCGTTCGTCAAGGATC |
| Consensus I-F Leader | EcoRI and BamHI digest of Addgene | AATTCGAGCTCGGTACCTCGCGAATGCATCTAGATCCATGGACCC TTTTTTCGGACGATTTAAAAAACCCTTATAAATCAGCAAGTTACGAG |

| | | |
|---|---|---|
| and consensus I-F Locus DNA | plasmid 192209, $^{32}$P-labelled for integration assays | ACCTCGAAAAAAAAGGGTTTCTGGCGGGAAAAACTCAAAAAACGC<br>TTCGAAATCAACCGGTTATAGGTTTTCGGAGCTAGTTCACTGCCGT<br>ATAGGCAGCTTAGAAAATCAGCCGGACGTTGTAGTAGTCGAGCGC<br>GGTGTTCACTGCCGTATAGGCAGCTTAGAAAGCCGGGTGGTGACT<br>TCTGGACGGCGTTCGTCAAGGATC |
| Consensus I-F leader and consensus I-E locus DNA | EcoRI and BamHI digest of Addgene plasmid 192210, $^{32}$P-labelled for integration assays | AATTCGAGCTCGGTACCTCGCGAATGCATCTAGATCCATGGACCC<br>TTTTTTCGGACGATTTAAAAAACCCTTATAAATCAGCAAGTTACGAG<br>ACCTCGAAAAAAAAGGGTTTCTGGCGGGAAAAACTCAAAAAACGC<br>TTCGAAATCAACCGGTTATAGGTTTTCGGAGCTAGTGTTCCCCGC<br>GCGAGCGGGGATGAACCGATCAGCCGGACGTTGTAGTAGTCGAG<br>CGCGGTGTGTTCCCCGCGCGAGCGGGGATGAACCGGCCGGGTG<br>GTGACTTCTGGACGGCGTTCGTCAAGGATC |
| Consensus I-F leader and consensus I-C locus DNA | EcoRI and BamHI digest of Addgene plasmid 192211, $^{32}$P-labelled for integration assays | AATTCGAGCTCGGTACCTCGCGAATGCATCTAGATCCATGGACCC<br>TTTTTTCGGACGATTTAAAAAACCCTTATAAATCAGCAAGTTACGAG<br>ACCTCGAAAAAAAAGGGTTTCTGGCGGGAAAAACTCAAAAAACGC<br>TTCGAAATCAACCGGTTATAGGTTTTCGGAGCTAGTCGCGCCCTT<br>CACGGGCGCGTGGATTGAAACATCAGCCGGACGTTGTAGTAGTC<br>GAGCGCGGTGTCGCGCCCTTCACGGGCGCGTGGATTGAAACGCC<br>GGGTGGTGACTTCTGGACGGCGTTCGTCAAGGATC |
| Consensus I-F leader and consensus II-A locus DNA | EcoRI and BamHI digest of Addgene plasmid 192212, $^{32}$P-labelled for integration assays | AATTCGAGCTCGGTACCTCGCGAATGCATCTAGATCCATGGACCC<br>TTTTTTCGGACGATTTAAAAAACCCTTATAAATCAGCAAGTTACGAG<br>ACCTCGAAAAAAAAGGGTTTCTGGCGGGAAAAACTCAAAAAACGC<br>TTCGAAATCAACCGGTTATAGGTTTTCGGAGCTAGTTTTAGAGTCA<br>TATTATTTAGAATGGTACCAAAACATCAGCCGGACGTTGTAGTAGT<br>CGAGCGCGGTGTTTTAGAGTCATATTATTTAGAATGGTACCAAAAC<br>GCCGGGTGGTGACTTCTGGACGGCGTTCGTCAAGGATC |
| Consensus I-E leader and consensus I-E locus DNA | EcoRI and BamHI digest of Addgene plasmid 192213, $^{32}$P-labelled for integration assays | AATTCGAGCTCGGTACCTCGCGAATGCATCTAGATCCATGGCGGT<br>GGACTTTGATGGCTTGAATCTGGTCGCCTTTCAGCCCTTGGAAAA<br>CCCTGATCTTTAACAAGGAAAAGTTGGTAGATTTTTAGCGGCTAAT<br>TTTCCCCTTTGGGGACAATTGGTTACGCTAAGAGTGTTCCCCGCG<br>CGAGCGGGGATGAACCGATCAGCCGGACGTTGTAGTAGTCGAGC<br>GCGGTGTGTTCCCCGCGCGAGCGGGGATGAACCGGCCGGGTGG<br>TGACTTCTGGACGGCGTTCGTCAAGGATC |
| Consensus I-C leader and consensus I-C locus DNA | EcoRI and BamHI digest of Addgene plasmid 192214, $^{32}$P-labelled for integration assays | AATTCGAGCTCGGTACCTCGCGAATGCATCTAGATCCATGGGAGT<br>GCGAACGTGTAGCGACCGGGATTTTACGGGGAGGTTCGCGGATT<br>TGCACGGCGTTGAATTTGTTAGGGATTTTTTTAAGCACGGTGGGAA<br>TTTGAGGACGGAGGCGTGTTTGGGCCACAGAAGAGCAGAGGTTC<br>GCGGAATTGGGCGGTTTTTGTCTGGCAGTACAGTGGGTTAGGTTA<br>GGTGGGTCGCGCCCTTCACGGGCGCGTGGATTGAAACATCAGCC<br>GGACGTTGTAGTAGTCGAGCGCGGTGTCGCGCCCTTCACGGGCG<br>CGTGGATTGAAACGCCGGGTGGTGACTTCTGGACGGCGTTCGTC<br>AAGGATC |
| Consensus II-A leader and consensus II-A locus DNA | EcoRI and BamHI digest of Addgene plasmid 192215, $^{32}$P-labelled for integration assays | AATTCGAGCTCGGTACCTCGCGAATGCATCTAGATCCATGGATCT<br>CTTTAAATAATGTGGATGTCCTGTTTTTAGAACAAAGGGTGGTCCA<br>GAACAGATTCCAATATATTTTGGACGAAAACTTTTATTTGAGTTATG<br>AAAAAGCTTAAATTGTTACTGATTAGTGGTTCATTCTAAACTGAAAT<br>CTAGCTATGGATAAGTGATGCGAGTACGGAACTTTGGAAAAAAATA<br>ATTCTCCGAGGTTTTAGAGTCATATTATTTAGAATGGTACCAAAACA<br>TCAGCCGGACGTTGTAGTAGTCGAGCGCGGTGTTTTAGAGTCATA<br>TTATTTAGAATGGTACCAAAACGCCGGGTGGTGACTTCTGGACGG<br>CGTTCGTCAAGGATC |
| I-F leader and I-F repeat construct not studied | EcoRI and BamHI digest of Addgene plasmid 192216, $^{32}$P-labelled for integration assays | AATTCGAGCTCGGTACCTCGCGAATGCATCTAGATCCATGGACCC<br>TTTTTTCGGACTTAAAAAACCCTTATAAATCAGCAAGTTACGAGAC<br>CTCGAAAAAAAAGGGTTTCTGGCGGGAAAAACTCAAAAAACGCTT<br>CGAAATCAACCGGTTATAGGTTTTCGGAGCTAGTTCACTGCCGTG<br>TAGGCAGCTAAGAAAATCAGCCGGACGTTGTAGTAGTCGAGCGCG |

| | | |
|---|---|---|
| further in this paper (X1) | | GTGTTCACTGCCGTGTAGGCAGCTAAGAAAGCCGGGTGGTGACTTCTGGACGGCGTTCGTCAAGGATC |
| I-F leader and I-F repeat construct not studied further in this paper (X2) | EcoRI and BamHI digest of Addgene plasmid 192217, $^{32}$P-labelled for integration assays | AATTCGAGCTCGGTACCTCGCGAATGCATCTAGATCCATGGACCCTTTTTTCGGACGATTTAAAAAACCCTTATAAATCAGCAAGTTACGAGACCTGATCGAAAAAAAAGGGTTTCTGGCGGAAAAACTCAAAAAACGCTTCGAAATCAACCGGTTATAGGTTTTCGGAGCTAGTTCACTGCCGTGTAGGCAGCTAAGAAAATCAGCCGGACGTTGTAGTAGTCGAGCGCGGTGTTCACTGCCGTGTAGGCAGCTAAGAAAGCCGGGTGGTGACTTCTGGACGGCGTTCGTCAAGGATC |
| I-F leader and I-F repeat construct with G1A,T2A mutation in repeat | EcoRI and BamHI digest of Addgene plasmid 200183, $^{32}$P-labelled for integration assays | GAATTCGAGCTCGGTACCTCGCGAATGCATCTAGATCCATGGACCCTTTTTTCGGACGATTTCTTACGCCCTTATAAATCAGCAAGTTACGAGACCTCGAAAAAAGAGGGTTTCTGGCGGAAAAACTCGGTATTTCTTTTTCCTTCAAATGGTTATAGGTTTTCGGAGCTAAATCACTGCCGTGTAGGCAGCTAAGAAAATCAGCCGGACGTTGTAGTAGTCGAGCGCGGTAATCACTGCCGTGTAGGCAGCTAAGAAAGCCGGGTGGTGACTTCTGGACGGCGTTCGTCAAGGATCC |
| I-F leader and I-F repeat construct with A28C mutation in the repeat | EcoRI and BamHI digest of Addgene plasmid 200184, $^{32}$P-labelled for integration assays | GAATTCGAGCTCGGTACCTCGCGAATGCATCTAGATCCATGGACCCTTTTTTCGGACGATTTCTTACGCCCTTATAAATCAGCAAGTTACGAGACCTCGAAAAAAGAGGGTTTCTGGCGGAAAAACTCGGTATTTCTTTTTCCTTCAAATGGTTATAGGTTTTCGGAGCTAGTTCACTGCCGTGTAGGCAGCTAAGAACATCAGCCGGACGTTGTAGTAGTCGAGCGCGGTGTTCACTGCCGTGTAGGCAGCTAAGAACGCCGGGTGGTGACTTCTGGACGGCGTTCGTCAAGGATCC |
| I-F leader and I-F repeat construct with G1A,T2A, A28C mutation in repeat | EcoRI and BamHI digest of Addgene plasmid 200185, $^{32}$P-labelled for integration assays | GAATTCGAGCTCGGTACCTCGCGAATGCATCTAGATCCATGGACCCTTTTTTCGGACGATTTCTTACGCCCTTATAAATCAGCAAGTTACGAGACCTCGAAAAAAGAGGGTTTCTGGCGGAAAAACTCGGTATTTCTTTTTCCTTCAAATGGTTATAGGTTTTCGGAGCTAAATCACTGCCGTGTAGGCAGCTAAGAACATCAGCCGGACGTTGTAGTAGTCGAGCGCGGTAATCACTGCCGTGTAGGCAGCTAAGAACGCCGGGTGGTGACTTCTGGACGGCGTTCGTCAAGGATCC |
| Cas1_H25A_mut_F | One of two primers pairs used to generate the Cas1$^{H25A}$ mutant | CTACCTGCAAgccTGCCGGGTACTG |
| Cas1_H25A_mut_R | One of two primers pairs used to generate the Cas1$^{H25A}$ mutant | TACAGGTTGGCACGCTTG |
| Cas1_E184A_mut_F | One of two primers pairs used to generate the Cas1$^{E184A}$ mutant | CCCAATCATGcGCATCTACTGACCG |
| Cas1_E184A_mut_R | One of two primers pairs used to generate the Cas1$^{E184A}$ mutant | AGCCTGCTCCAACGCTCG |
| Cas2/3_K11D,R12E_mut_F | One of two primers pairs used to generate the Cas2/3$^{K11D,R12E}$ mutant | GCAATGCGAAgatgaaGCCCTGAGCGAAAC |

| | | |
|---|---|---|
| Cas2/3_K11D,R12E_mut_R | One of two primers pairs used to generate the Cas2/3$^{K11D,R12E}$ mutant | GACACCAGCAGGATGTTC |
| Cas2/3_R55E,N56D_mut_F | One of two primers pairs used to generate the Cas2/3$^{R55E,N56D}$ mutant | AAGCGCACGGgaggacACCGCCGTAG |
| Cas2/3_R55E,N56D_mut_R | One of two primers pairs used to generate the Cas2/3$^{R55E,N56D}$ mutant | TTCTTCAGCAGGCGTCGC |
